# Lysosome positioning affects NLRP3 inflammasome activation

**DOI:** 10.1101/2025.03.21.644552

**Authors:** Billie J. Matchett, Jack P. Green, Paula I. Seoane, Abigail Owen, Tomasz Drobkiewicz, Hayley Bennett, Antony Adamson, Gloria Lopez-Castejon, Christopher Hoyle, Martin Lowe, David Brough

## Abstract

The NLRP3 inflammasome, an important multi-molecular protein complex driving inflammation, is activated by multiple cellular stresses, including a loss of organelle homeostasis. Studies indicate that lysosome disruption can play a role in NLRP3 inflammasome activation, but whether lysosome positioning impacts NLRP3 inflammasome activation is currently unknown. Using HeLa and macrophage cell models, the effect of repositioning lysosomes to either the cell periphery or the perinuclear region on inflammasome activation was studied. Lysosome positioning to the cell periphery increased NLRP3 inflammasome activation in response to nigericin. This effect correlated with a change in the subcellular localisation of the inflammasome complex and an increase in its size. As macrophages at sites of tissue inflammation are exposed to environments known to influence lysosome position, these data help us understand how subcellular organelle reorganisation can regulate inflammation in response to inflammatory stimuli.

## Introduction

Inflammation is regulated by our immune system, occurring in response to pathogens and tissue injury to promote healing (Chen et al., 2018). An inflammatory response is triggered following detection of harmful stimuli that induce cellular signalling, leading to the release of inflammatory cytokines and recruitment of immune cells (Chen et al., 2018). Despite being important for maintaining health, inflammation also contributes to the worsening of many diseases, including stroke, psoriasis, rheumatoid arthritis and inflammatory bowel disease (Amor et al., 2010, Scotece and Conde-Aranda, 2022).

Inflammasomes are early initiators of inflammatory responses. Inflammasomes are formed following the activation of specific cytoplasmic pattern recognition receptors (PRRs). Inflammasome-forming PRRs include members of the NOD-like receptor (NLR) family, such as NLRP1, NLRP3, NLRP6 or NLRC4, and PYHIN protein family members such as absent in melanoma (AIM2) (Schroder and Tschopp, 2010, Zheng et al., 2020). The best characterised inflammasome is formed following activation of the PRR NLRP3, which triggers recruitment of the adaptor protein apoptosis-associated speck-like protein containing a CARD (ASC) (Schroder and Tschopp, 2010, Zheng et al., 2020). The ASC scaffold recruits and activates caspase-1, which subsequently promotes the proteolytic cleavage of the pro-inflammatory cytokine precursors pro-IL-1β and pro-IL-18 to their active forms (Chan and Schroder, 2020). Caspase-1 also cleaves gasdermin D, a pore-forming protein that allows cytokine release and subsequently triggers NINJ1-dependent cell lysis (Aglietti et al., 2016, Degen et al., 2023, He et al., 2015, Kayagaki et al., 2021, Liu et al., 2016, Sborgi et al., 2016).

In macrophages, canonical activation of the NLRP3 inflammasome occurs via a two-step process that first involves a priming signal, typically lipopolysaccharide (LPS)-induced Toll-like receptor 4 (TLR4) activation, that induces NLRP3 and pro-IL-1β expression (Bauernfeind et al., 2009, Franchi et al., 2009). The second step is an activation signal, which occurs in response to a range of pathogen-or damage-associated molecular patterns (PAMPs and DAMPs, respectively), such as the K^+^ ionophore nigericin, extracellular ATP (Mariathasan et al., 2006), silica crystals (Hornung et al., 2008), and imiquimod (Gross et al., 2016). These structurally diverse PAMPs and DAMPs may activate the NLRP3 inflammasome by altering cellular homeostasis (Liston and Masters, 2017), such as causing organelle stress and dysfunction (Seoane et al., 2020). Previous studies suggest that the endolysosomal system is important for NLRP3 inflammasome activation. For example, NLRP3-activating stimuli disrupt endolysosomal cargo trafficking, resulting in the accumulation of phosphatidylinositol 4-phosphate (PI4P) in endolysosomal membranes, facilitating NLRP3 recruitment (Chen and Chen, 2018, Lee et al., 2023, Zhang et al., 2023). Phagocytosis of particulates such as silica, or aluminum salt crystals, and the lysosome membrane-disrupting chemical L-leucyl-L-leucine methyl ester (LLOMe), induce lysosome membrane permeablilisation and rupture, leading to NLRP3 inflammasome activation (Hornung et al., 2008, Lima et al., 2013) via K^+^ efflux, an established trigger of NLRP3 inflammasome activation (Munoz-Planillo et al., 2013). Spatiotemporal proteomic profiling shows significant changes in the organellar proteomes of endosomes and lysosomes after stimulation of macrophages with the NLRP3-activating stimuli nigericin and CL097 (Hollingsworth et al., 2024). Furthermore, the lysosomal complex Ragulator is also reported to be involved in NLRP3 inflammasome activation by interacting with NLRP3 directly, with reports of ASC specks forming near lysosomes (Tsujimoto et al., 2023). Together, this evidence suggests that endosomes and lysosomes are heavily involved in the activation of the NLRP3 inflammasome.

Lysosomes regulate many cellular functions, including degradation and recycling of cellular waste, cellular signalling and metabolism, cell death and inflammation (Settembre and Perera, 2024). Although mostly concentrated in the perinuclear region surrounding the microtubule-organising centre (MTOC) (Jongsma et al., 2016), lysosomes are highly dynamic organelles, trafficking throughout the cell via microtubules and motor proteins (Pu et al., 2016). Trafficking of lysosomes towards the cell periphery (plus-end or anterograde transport) is mediated by kinesin motors such as kinesin-1 (Hollenbeck and Swanson, 1990), alongside adaptor proteins such as BLOC-1-related complex (BORC), the Arf-like small GTPase Arl8 (Arl8a/Arl8b) and SKIP (Bagshaw et al., 2006, Hofmann and Munro, 2006, Pu et al., 2015, Rosa-Ferreira and Munro, 2011). Conversely, trafficking of lysosomes towards the perinuclear region (minus-end or retrograde transport) is mediated by dynein-dynactin motors (Burkhardt et al., 1997, Lin and Collins, 1992), alongside adaptor proteins such as Rab7, Rab-interacting lysosomal protein (RILP), JIP4 and TMEM55B (Cantalupo et al., 2001, Willett et al., 2017).

The role of lysosome positioning in the regulation of different cellular processes is becoming increasingly apparent. For example, starvation-induced autophagy causes lysosomes to concentrate around the perinuclear region, which favours autophagosome formation and autophagosome–lysosome fusion (Willett et al., 2017). Conversely, high nutrient levels induce repositioning of the lysosomes to the cell periphery, increasing mTOR activity, blocking autophagosome synthesis and reducing autophagosome–lysosome fusion (Korolchuk et al., 2011). Recently, lysosome positioning has been implicated in inflammation. LPS and IFNγ treatment causes lysosome movement in macrophages (Char et al., 2023), and lysosome positioning is implicated in TLR7 signalling, turnover and trafficking in the context of human lupus (Mishra et al., 2024). However, whether lysosomal positioning regulates NLRP3 inflammasome activation is currently unknown.

Given the role of lysosomes in NLRP3 inflammasome activation, and the emerging importance of lysosome position for lysosome function, the aim of this study was to investigate the influence of lysosome position on NLRP3 inflammasome activation. Using HeLa and macrophage cell models, we show that manipulating lysosome positioning disrupted endolysosomal trafficking, induced lysosomal damage and altered lysosomal pH. Inducing peripheral lysosome positioning increased NLRP3 inflammasome activation in response to nigericin. Manipulating lysosome positioning in macrophages influenced the subcellular localisation and the size of the ASC speck, consistent with lysosome position influencing inflammasome assembly. Together, these data suggest that lysosome positioning plays a role in NLRP3 inflammasome activation, and could represent a new avenue for therapeutic intervention.

## Results

### Arl8b-mCherry and TMEM55B-mCherry overexpression in HeLa cells alters lysosome position

Initially, we set out to establish an experimental set up where we could reliably manipulate the position of lysosomes within the cell, which is dependent upon the microtubule network, as well as the motor proteins and specific adaptor proteins that tether them (Cabukusta and Neefjes, 2018). To drive lysosome position to the cell periphery, we overexpressed Arl8b, a lysosomal GTPase that binds to the microtubule motor kinesin-1 (Johnson et al., 2016, Rosa-Ferreira and Munro, 2011). To move lysosomes to the perinuclear space, we overexpressed TMEM55B, an adaptor protein that connects lysosomes to dynein-dynactin motors (Willett et al., 2017). We generated stable HeLa cell-lines expressing either the fluorescent protein mCherry alone for our control cell line, Arl8b-mCherry, or TMEM55B-mCherry (Fig S1A, B). HeLa cells were labelled for both the lysosome marker LAMP1 and the microtubule marker α-tubulin and imaged using fluorescence microscopy (Fig 1A and B). mCherry fluorescent cells were specifically selected for analysis and lysosome position was assessed using the ImageJ plugin RadialIntensity profile as reported by Filipek *et al*. and suggested as a principal method of lysosome position analysis (Barral et al., 2022, Filipek et al., 2017) (Fig S1C). This analysis showed that Arl8b overexpression shifted the distribution of lysosomes towards the cell periphery (Fig 1C and E) and that TMEM55B overexpression caused the lysosome distribution to collapse toward the perinuclear region (Fig 1D and F). The cells used were mixed pools and not clonal populations, and thus heterogeneity in expression levels of transduced components would be expected, although this did not prevent us from observing significant shifts in lysosome position. To examine the effect of Arl8b and TMEM55B overexpression on other endo-lysosomal compartments, we labelled early endosomes using EEA1 (Fig S2A and B). These data show that Arl8b or TMEM55B overexpression had no significant effect upon the distribution of early endosomes (Fig S2). Together, these data suggest that overexpression of Arl8b and TMEM55B preferentially alters lysosome distribution in HeLa cells, in line with previous studies (Johnson et al., 2016, Willett et al., 2017).

**Figure 1.**
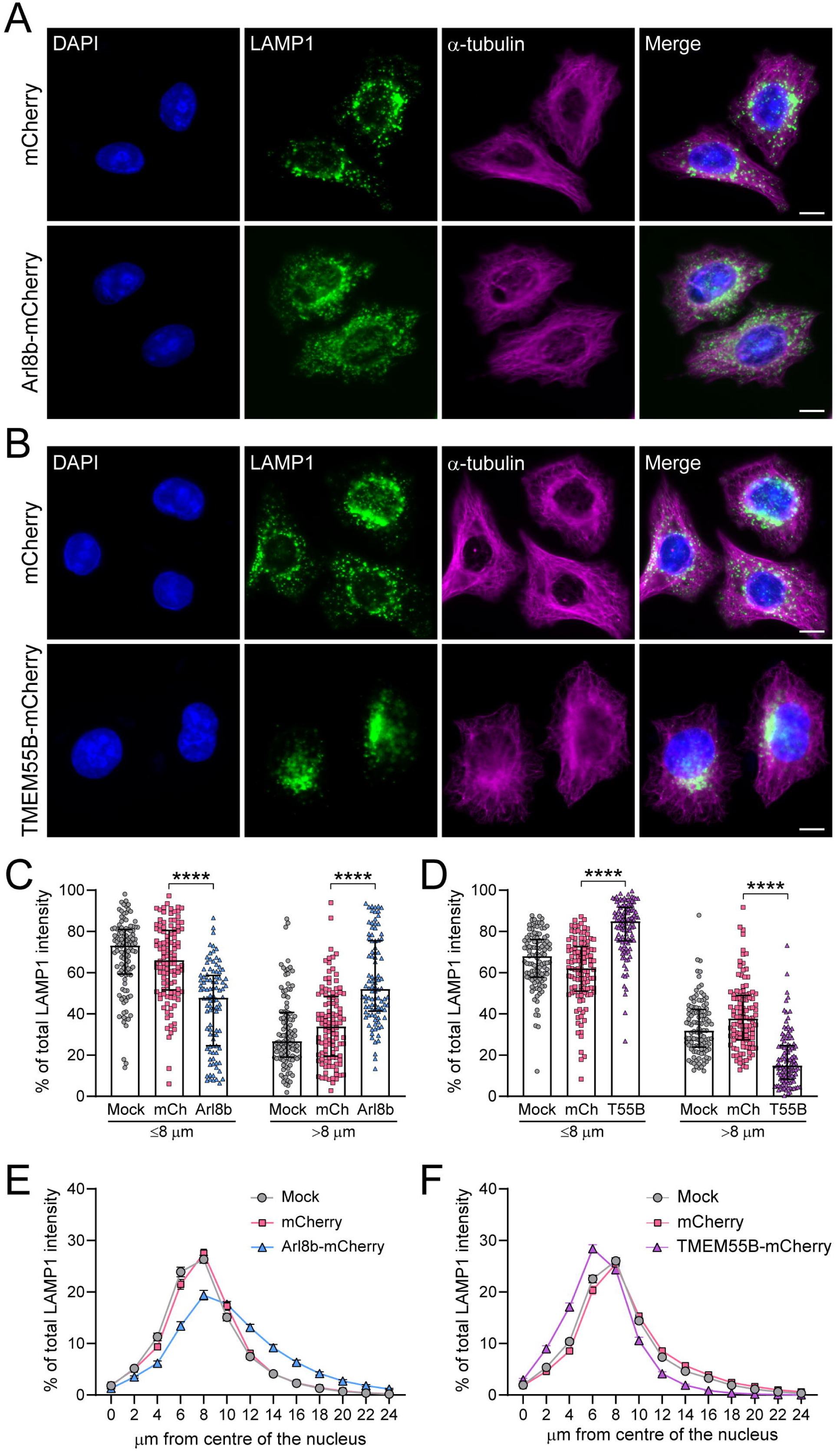
Arl8b-mCherry and TMEM55B-mCherry overexpression in HeLa cells positions lysosomes to the periphery and perinuclear regions, respectively. **[A and B]** Representative images for HeLa mCherry, alongside HeLa Arl8b-mCherry **[A]** and HeLa TMEM55B-mCherry **[B]**, stained for the lysosomal marker LAMP1 (green) and the microtubule marker α-tubulin (magenta). Scale bar = 10 μm. Images are representative of 36 random fields of view per cell line from three independent experiments. **[C and D]** Quantification of the percentage total LAMP1 intensity at ≤ 8 µm (perinuclear localisation) and > 8 µm (peripheral localisation) from the centre of the nucleus in HeLa Arl8b-mCherry [C] and HeLa TMEM55B-mCherry cells **[D]**. *n* = 110 (UT), 110 (mCherry), 96 (Arl8b-mCherry), 110 (TMEM55B-mCherry) from three independent repeats. **[E and F]** Quantification of the percentage total LAMP1 intensity at each 2 μm interval in HeLa Arl8b-mCherry **[E]** and HeLa TMEM55B-mCherry **[F]** cell lines, showing lysosome distribution within the cell. *n* = 110 (UT), 110 (mCherry), 96 (Arl8b-mCherry), 110 (TMEM55B-mCherry) from three independent repeats. Data were analysed using Kruskal-Wallis test with Dunn’s multiple comparison test. ****P ≤ 0.0001.

### Altering lysosome position affects endo-lysosomal cargo trafficking in HeLa cells

We next sought to characterise the effect of manipulating lysosome position on subcellular trafficking. Trafficking from endosomes to lysosomes was assessed using an established epidermal growth factor (EGF) trafficking assay (Liang et al., 2008, Noakes et al., 2011). This assay tracks fluorescently labelled EGF from the cell surface via endosomes to the lysosome, where it is ultimately degraded. This degradation is reflected in a reduction in EGF fluorescence, and therefore defects in this pathway are indicated through maintained or increased EGF fluorescence. Our previous study reported defects in this pathway in response to NLRP3 inflammasome-activating stimuli (Lee et al., 2023). Therefore, to test whether altered lysosome position affects endo-lysosomal trafficking, cells were incubated in media containing fluorescently-labelled EGF and images were taken at 0, 1 and 2 h time points to assess changes in EGF fluorescence (Fig 2A). EGF trafficking to the lysosome was reduced in Arl8b-overexpressing cells at 1 h, and in both Arl8b- and TMEM55B-overexpressing cells at 2 h post incubation (Fig 2B), suggesting lysosome position affects endosome-to-lysosome cargo trafficking.

**Figure 2.**
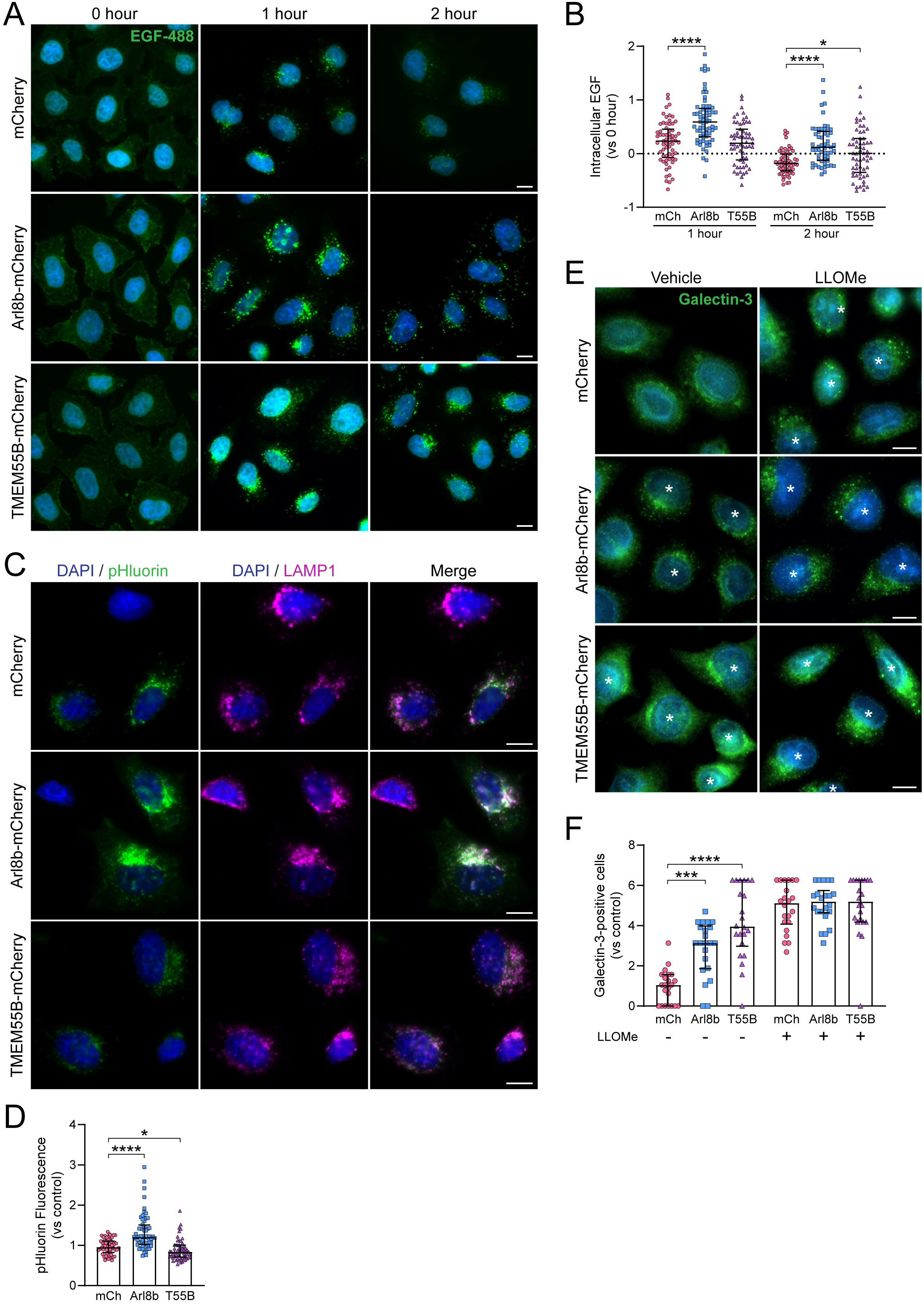
Manipulated lysosomal positioning disrupts endo-lysosomal trafficking, alters lysosome luminal pH and damages lysosomes in HeLa cells. **[A]** Fluorescence images of EGF-488 (green) in all three HeLa cell lines. Cells were incubated with 0.4 µg mL^−1^ EGF Alexa Fluor 488 for 1 h on ice, followed by incubation in media at 37°C for 0, 1 and 2 h. Scale bar = 10 μm. Images are representative of 18 random fields of view per cell line per timepoint from three independent experiments. **[B]** Quantification of EGF-488 degradation in mCherry, Arl8b-mCherry and TMEM55B-mCherry HeLa cells was measured by the fluorescence intensity of EGF-488 at 1 and 2 h after incubation. Data are presented as normalised to EGF-488 fluorescence intensity at 0 hour. *n* = 65 (mCherry, 1 h), 59 (mCherry, 2 h), 65 (Arl8b-mCherry, 1 h), 57 (Arl8b-mCherry, 2 h), 59 (TMEM55B-mCherry, 1 h), 59 (TMEM55B-mCherry, 2 h) from 18 random fields of view from three independent experiments. **[C]** Representative images of pHluorin fluorescence (green) and LAMP1 (magenta) in all three HeLa cell lines. Scale bar = 10 μm. Images are representative of 24 random fields of view per cell line from four independent experiments. **[D]** Quantification of pHluorin fluorescence in all three HeLa cell lines. Data are presented as pHluorin fluorescence intensity normalised to mCherry control cell line. *n* = 62 (mCherry), 60 (Arl8b-mCherry) and 59 (TMEM55B-mCherry) from 24 random fields of view from four independent experiments. **[E]** Representative fluorescent images of galectin-3 (green) in all three HeLa cell lines following incubation with vehicle (ethanol; 0.5% v/v) or the lysosome-destabilising agent LLOMe (1 mM) for 30 min. Asterisks denote cells positive for galectin-3 puncta. Scale bar = 10 μm. Images are representative of 22 random fields of view per cell line per treatment from four independent experiments. **[F]** Quantification of cells positive for galectin-3 puncta following 30 min stimulation with vehicle or LLOMe in all three HeLa cell lines. Data are presented as normalised to vehicle-treated mCherry control cell line. *n* = 22 random fields of view per cell line per treatment from four independent experiments. Data were analysed using Kruskal-Wallis test with Dunn’s multiple comparison test. *P ≤ 0.05, ***P ≤ 0.001, ****P ≤ 0.0001.

### Altering lysosome position affects lysosome pH and integrity in HeLa cells

Next, we investigated the role of lysosome positioning on lysosome pH and integrity. First, we analysed lysosomal pH in our cell lines using the lysosome-specific pH sensor Lyso-pHluorin (Rost et al., 2015) (Fig 2C). In the acidic environment of the lysosome, Lyso-pHluorin is non-fluorescent, but as the lysosome becomes more alkaline, which occurs during lysosomal damage, Lyso-pHluorin becomes fluorescent, enabling us to examine changes in lysosomal pH (Miesenbock et al., 1998, Rost et al., 2015). We observed that Arl8b-overexpressing cells had lysosomes with a more alkaline pH at baseline, whereas TMEM55B-overexpressing cells had a slightly more acidic pH at baseline (Fig 2D). These results are consistent with previous reports showing that lysosomes positioned in the cell periphery have more alkaline luminal pH, in contrast to perinuclear lysosomes which exhibit a more acidic pH (Johnson et al., 2016).

We also tested whether manipulating lysosome position affects lysosomal stress and damage by assessing galectin-3. Galectin-3 is a cytosolic protein that, upon lysosomal damage, accumulates on the surface of damaged lysosomes where it forms puncta (Aits et al., 2015). Therefore, cells with lysosomal damage exhibit galectin-3-positive puncta. To assess this, Arl8b- and TMEM55B-overexpressing cells were stained for galectin-3, and the percentage of cells with galectin-3-positive puncta was assessed (Fig 2E, see asterisks). The lysosome-disrupting agent LLOMe (Maejima et al., 2013) was used as a positive control. An increase in cells with galectin-3-positive puncta was observed at baseline in both Arl8b- and TMEM55B-overexpressing cells (Fig 2F), suggesting that manipulation of lysosome positioning caused lysosomal stress. No difference in the extent of lysosomal damage was observed between the cell lines following treatment with LLOMe, suggesting that these changes are only apparent under basal conditions.

### Peripheral lysosome positioning causes lysosome damage in macrophages

Next, we made a stable immortalised bone marrow-derived macrophage (iBMDM) cell-line with Arl8b-mCherry to study the effects of peripheral lysosome positioning on inflammatory processes in macrophages. Overexpression of TMEM55B in the iBMDMs was not well tolerated and the cells were not viable (data not shown). Arl8b-mCherry expressing iBMDMs were stained for LAMP1 and α-tubulin and imaged using fluorescence microscopy (Fig 3A). We observed that, similar to HeLa cells, Arl8b overexpression in iBMDMs shifted the distribution of lysosomes towards the cell periphery (Fig 3A-C), suggesting that this is a suitable model to assess the effects of peripheral lysosome positioning on inflammatory processes. We then investigated whether peripheral lysosome positioning in iBMDMs affects lysosomal integrity by assessing galectin-3 staining (Fig 3D). Arl8b overexpression increased the percentage of cells with galectin-3-positive puncta at baseline, with a further increase seen in LLOMe-treated cells (Fig 3D and E). This indicates that, as seen in HeLa cells, peripheral lysosome positioning caused lysosomal stress in macrophages.

**Figure 3.**
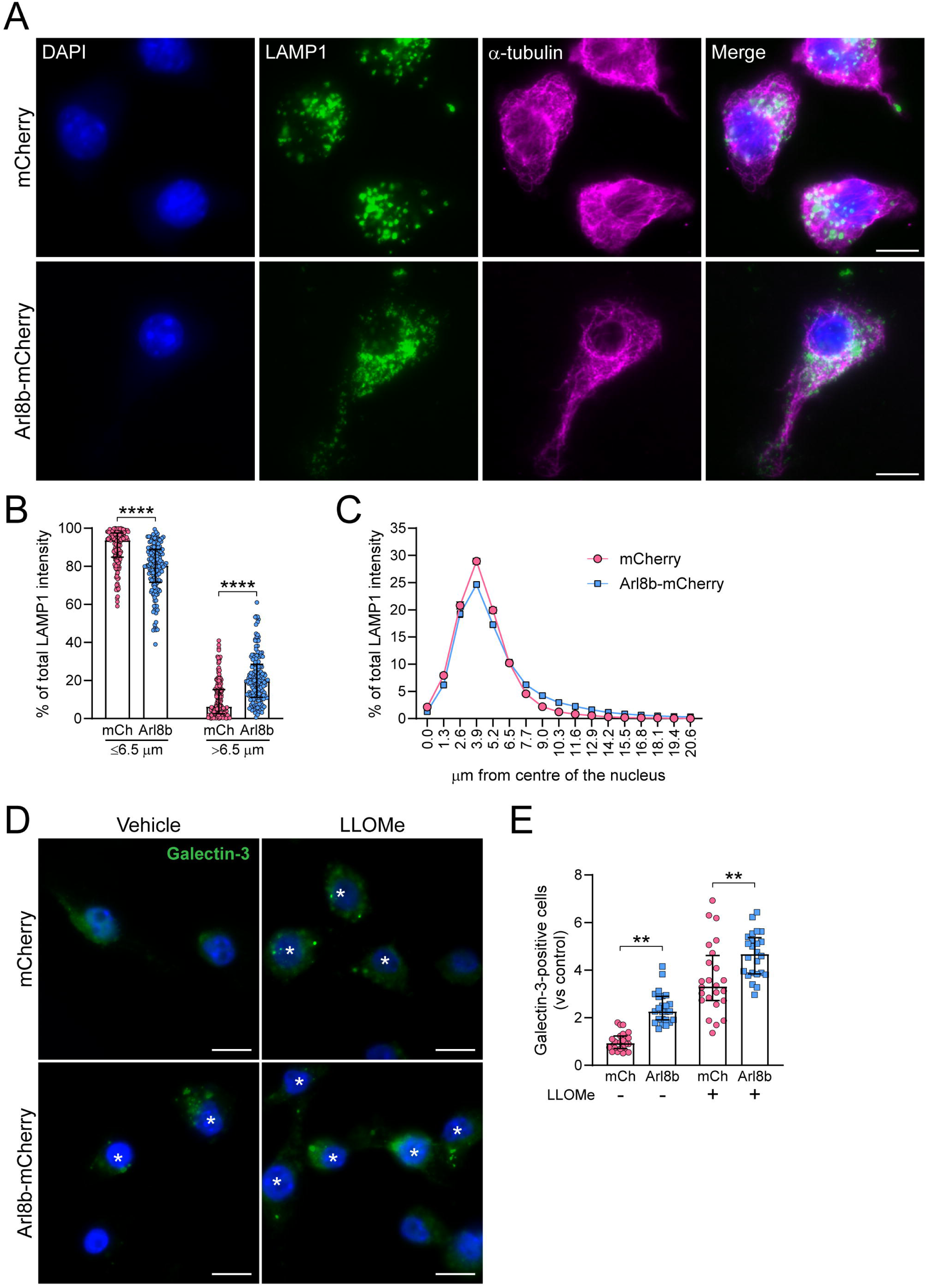
Arl8b-mCherry overexpression redistributes lysosomes to the periphery in iBMDMs and causes lysosomal damage. **[A-E]** iBMDM mCherry and iBMDM Arl8b-mCherry cells were primed with LPS (1 µg mL^−1^) in complete media for 2 h. **[A]** Representative images for each LPS-primed iBMDM cell line stained for the lysosomal marker LAMP1 (green), and the microtubule marker α-tubulin (magenta). Scale bar = 10 μm. Images are representative of 72 random fields of view per cell line from four independent experiments. **[B]** Quantification of the percentage total LAMP1 fluorescence intensity at ≤ 6.5 µm (perinuclear localisation) and > 6.5 µm (peripheral localisation) from the centre of the nucleus in each iBMDM cell line (*n* = 159 individual cells per cell line from four independent repeats). **[C]** Quantification of the percentage total LAMP1 fluorescence intensity at each 1.3 μm interval in iBMDMs expressing mCherry or Arl8b-mCherry, showing lysosome distribution within the cell (*n* = 159 individual cells per cell line from four independent repeats). **[D]** Representative fluorescence images of galectin-3 (green) in both iBMDM cell lines. LPS-primed iBMDMs were incubated with vehicle (ethanol; 0.5% v/v) or the lysosome-destabilising agent LLOMe (1 mM) for 30 min. Asterisks denote cells positive for galectin-3 puncta. Scale bar = 10 μm. Images are representative of 24 random fields of view per cell line per treatment from four independent experiments. **[E]** Quantification of cells positive for galectin-3 puncta, a measure of lysosomal damage, following 30 min stimulation with vehicle or LLOMe in both iBMDM cell lines. Data are presented as normalised to vehicle-treated mCherry control cell line. (*n* = 24 random fields of view per cell line per treatment from four independent experiments). Data were analysed using unpaired two-tailed Mann-Whitney test. **P ≤ 0.01, ****P ≤ 0.0001.

### Peripheral lysosome positioning does not affect priming but increases activation of the NLRP3 inflammasome to nigericin

To assess the relationship between lysosome position and NLRP3 inflammasome activation, we first assessed whether peripheral lysosome positioning affects NLRP3 inflammasome priming by stimulating iBMDM cells with LPS for 4 h. We observed an increase in the expression of the *Casp1* gene using qPCR in Arl8b-overexpressing cells (Fig 4Ai), however there was no significant difference in the relative gene expression of *Nlrp3* (Fig 4Aii) or *Pycard* (ASC, Fig 4Aiii), nor of pro-inflammatory cytokine genes (Fig 4Aiv-vi). Upon western blot analysis, we observed no difference in the levels of NLRP3 inflammasome components at baseline or following LPS priming between the two cell lines, although pro-IL-1β was slightly reduced in Arl8b-overexpressing iBMDMs (Fig 4B). There was no change in TNF-α (Fig 4C) or IL-6 (Fig 4D) secretion following LPS treatment between control and Arl8b-overexpressing iBMDMs. Together, these data suggest that peripheral lysosome positioning did not affect priming of the NLRP3 inflammasome.

**Figure 4.**
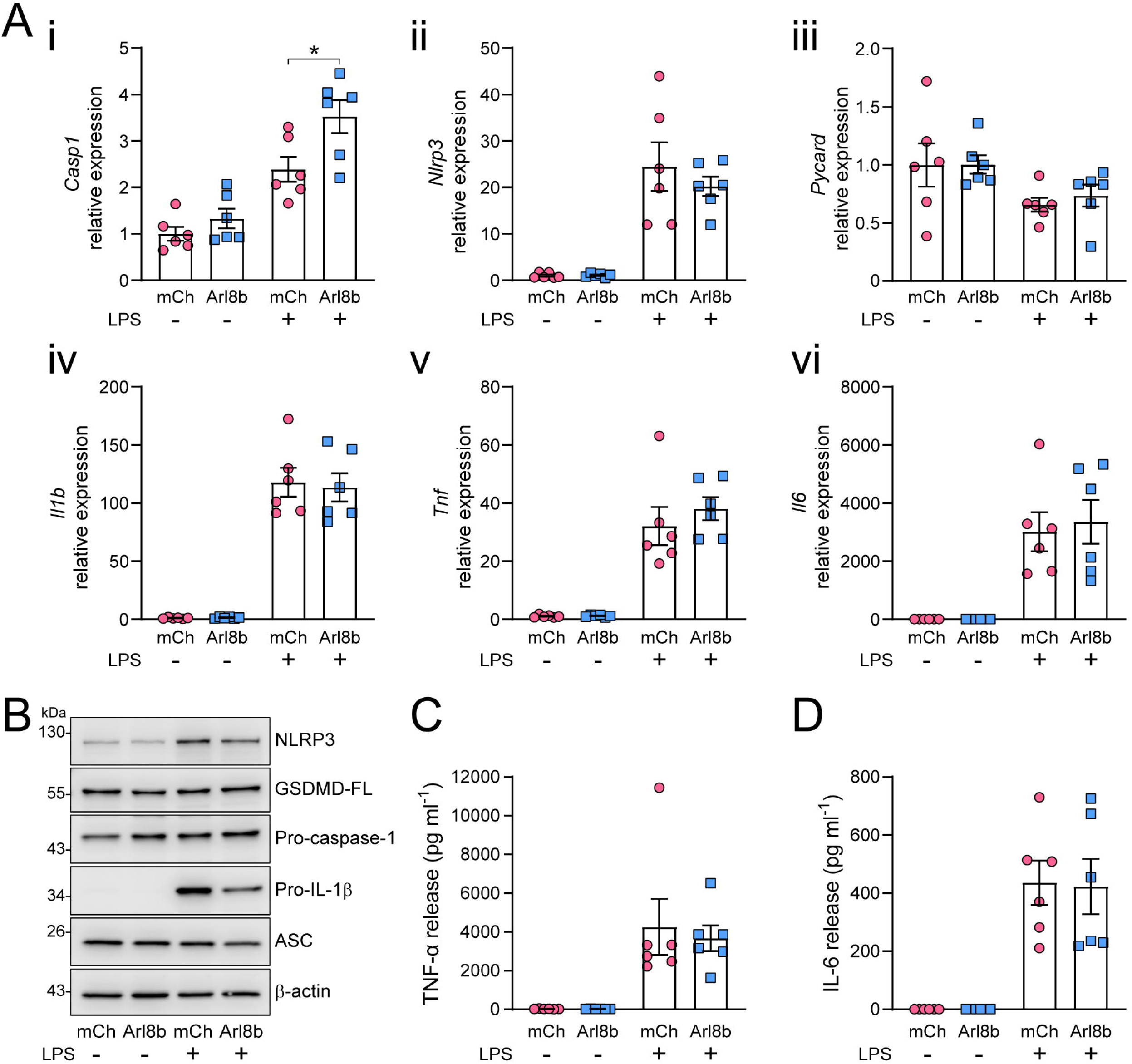
Effects of lysosome positioning on NLRP3 inflammasome priming. **[A-D]** iBMDM mCherry and iBMDM Arl8b-mCherry cells were primed with vehicle or LPS (1 µg mL^−1^) in complete media for 4 h. **[A]** qPCR analysis of TLR4-dependent genes and NLRP3 inflammasome components, with relative expression to the housekeeping gene *Hmbs* normalised to vehicle-treated mCherry cells. Genes measured are i) *Casp1* ii) *Nlrp3* iii) *Pycard* iv) *Il1b* v) *Tnf* vi) *Il6*. **[B]** Western blot of cell lysates showing expression of NLRP3 inflammasome components, showing NLRP3, full-length gasdermin D, pro-caspase-1, pro-IL-1β and ASC. Western blot is representative of 4 independent experiments. **[C and D]** Quantification of TNF-α **[C]** and IL-6 **[D]** release into the cell supernatant (measured by ELISA) (*n* = 6 independent experiments). Data were analysed using unpaired two-tailed Mann-Whitney test. *P ≤ 0.05.

We next assessed whether peripheral lysosome positioning affects NLRP3 inflammasome activation. LPS-primed Arl8b-overexpressing iBMDMs had increased cell death (Fig 5A), caspase-1 activity (Fig 5B) and IL-1β release (Fig 5C) following stimulation with nigericin. Each effect was NLRP3-dependent as the enhancement was blocked by the inhibitor MCC950 (Fig 5A-C), suggesting that peripheral lysosome positioning increases NLRP3 inflammasome activation in response to nigericin. To investigate whether increased NLRP3 inflammasome activation was due to an increase in the number of cells that formed an ASC speck, we next stimulated the iBMDMs with nigericin and stained for ASC specks (Fig 5D). There was no significant difference in the percentage of cells with ASC specks between mCherry or Arl8b-mCherry expressing iBMDMs (Fig 5E), suggesting that the increases observed in Fig 5A-C were not because of an increased number of inflammasomes being formed.

**Figure 5.**
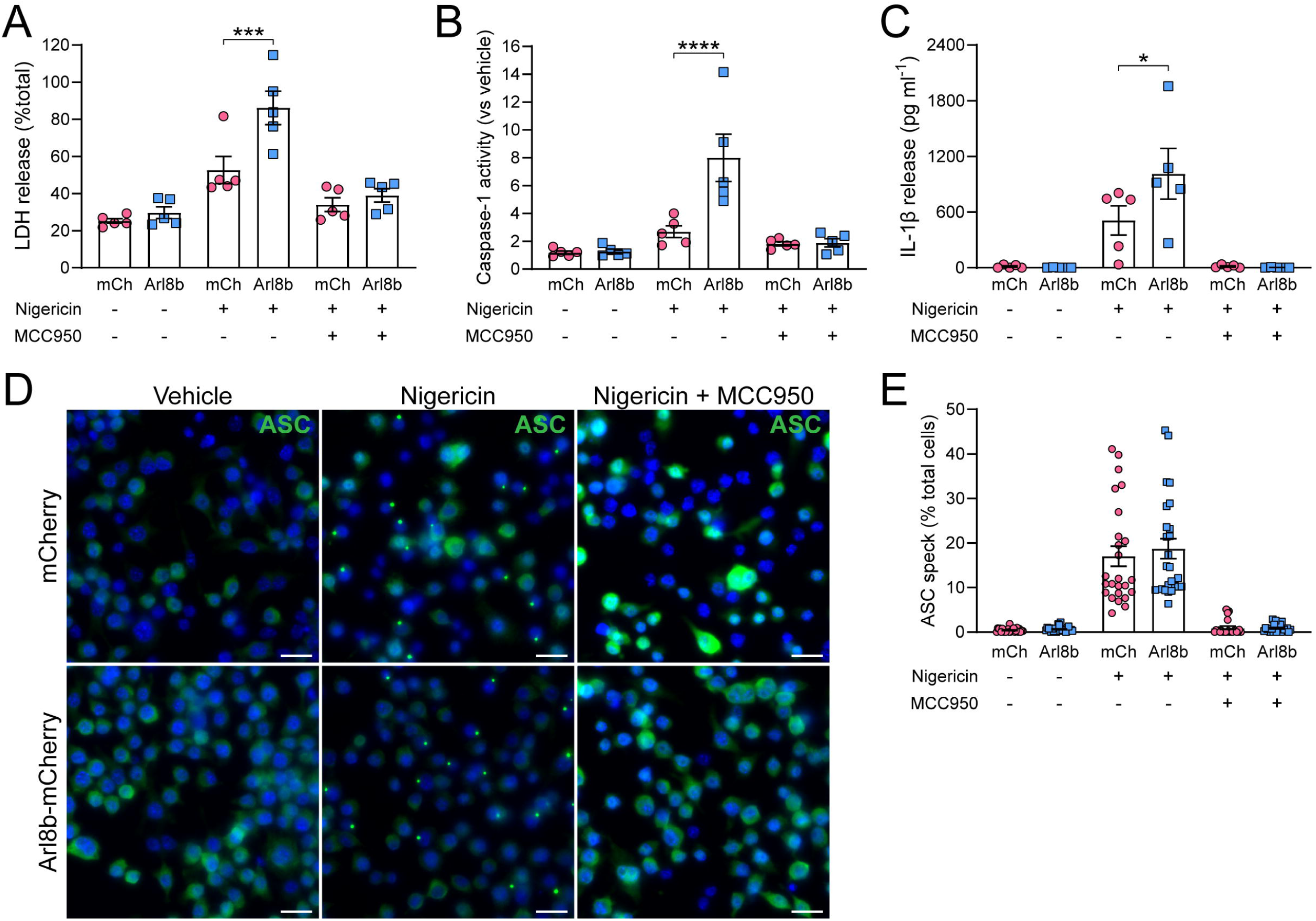
Peripheral lysosome positioning increases NLRP3 inflammasome activation in response to nigericin in iBMDMs. **[A-E]** iBMDM mCherry and iBMDM Arl8b-mCherry cells were primed with LPS (1 µg mL^−1^) in complete media for 2 h. **[A-C]** LPS-primed iBMDMs were pre-treated with PBS or MCC950 (10 µM) for 15 min before stimulation with vehicle or nigericin (10 µM) for 1 h. Supernatants were assessed for cell death (LDH release) **[A]**, caspase-1 activity (measured by Caspase-1 Glo) **[B],** and IL-1β release (measured by ELISA) **[C]** (*n* = 5 independent experiments). **[D and E]** LPS-primed iBMDMs were incubated with VX765 (10 µM) with or without MCC950 (10 µM) for 15 min before stimulation with vehicle (ethanol; 0.5% v/v) or nigericin (10 µM) for 1 h. **[D]** Representative images of ASC specks (green) in each iBMDM cell line treated with either vehicle, nigericin, or nigericin and MCC950. Scale bar = 20 µm. Images are representative of 25 random fields of view per cell line per treatment from five independent experiments. **[E]** Quantification of the percentage of cells positive for an ASC speck in both cell lines following stimulation (*n* = 25 random fields of view per cell line per treatment from five independent experiments). Data were analysed using a two-way ANOVA with Holm-Sidak’s post hoc test [A-C] or an unpaired two-tailed Mann-Whitney test [E]. *P ≤ 0.05, ***P ≤ 0.001, ****P ≤ 0.0001.

We next evaluated the effect of peripheral lysosome positioning on NLRP3 inflammasome activation in response to the NLRP3 activators silica, ATP and imiquimod. There were no significant differences in cell death or caspase-1 activity in response to silica, ATP, or imiquimod in iBMDMs (Fig S3A-I). However, despite there being no change in cell death (Fig S3D), there was a significant decrease in IL-1β release in response to ATP in Arl8b-overexpressing iBMDMs (Fig S3F). Interestingly, all stimuli showed a trend of decreased IL-1β release in Arl8b-overexpressing cells (Fig S3C, F and I), in contrast to the effect observed upon nigericin stimulation (Fig 5C). To determine whether the observed effects were specific to NLRP3, we also investigated the effects of peripheral lysosome positioning on activation of the AIM2 inflammasome. To do this, LPS-primed iBMDMs were transfected with the synthetic double-stranded DNA sequence poly(dA:dT) (Fernandes-Alnemri et al., 2009). To ensure any effects were AIM2-dependent, a control group was included where the cells were pre-incubated with 4-sulfonic calix[8]arene, an AIM2 inflammasome inhibitor (Green et al., 2023). No differences in cell death, caspase-1 activity, or IL-1β release were observed in response to poly(dA:dT) (Fig S3J-L), suggesting that peripheral lysosome positioning did not influence AIM2 inflammasome activation.

### Effect of lysosome position on ASC speck size and localization

We next assessed whether lysosome position affects the localisation and size of the ASC speck. HeLa cells do not express inflammasome components such as NLRP3 or ASC, so these were ectopically expressed by transfection. We co-transfected mCherry, Arl8b-mCherry, and TMEM55B-mCherry HeLa cell lines with wild-type mVenus-tagged NLRP3 (NLRP3-mVenus-WT) and full length ASC, and stained them with an ASC antibody (Fig 6A). We then assessed the number of NLRP3-ASC specks, the distance of the NLRP3-ASC speck from the centre of the nucleus, and the diameter of the NLRP3-ASC speck. There was no significant difference in the number of cells positive for a NLRP3-ASC speck between the three cell lines, suggesting that manipulating lysosome position does not play a role in the ability of a cell to form an inflammasome (Fig 6B). Furthermore, we observed no significant difference in the distance of the NLRP3-ASC speck from the centre of the nucleus (Fig 6C), nor the NLRP3-ASC speck diameter (Fig 6D), suggesting that lysosome positioning does not regulate subcellular location nor the size of the NLRP3-ASC speck in this experimental model.

**Figure 6.**
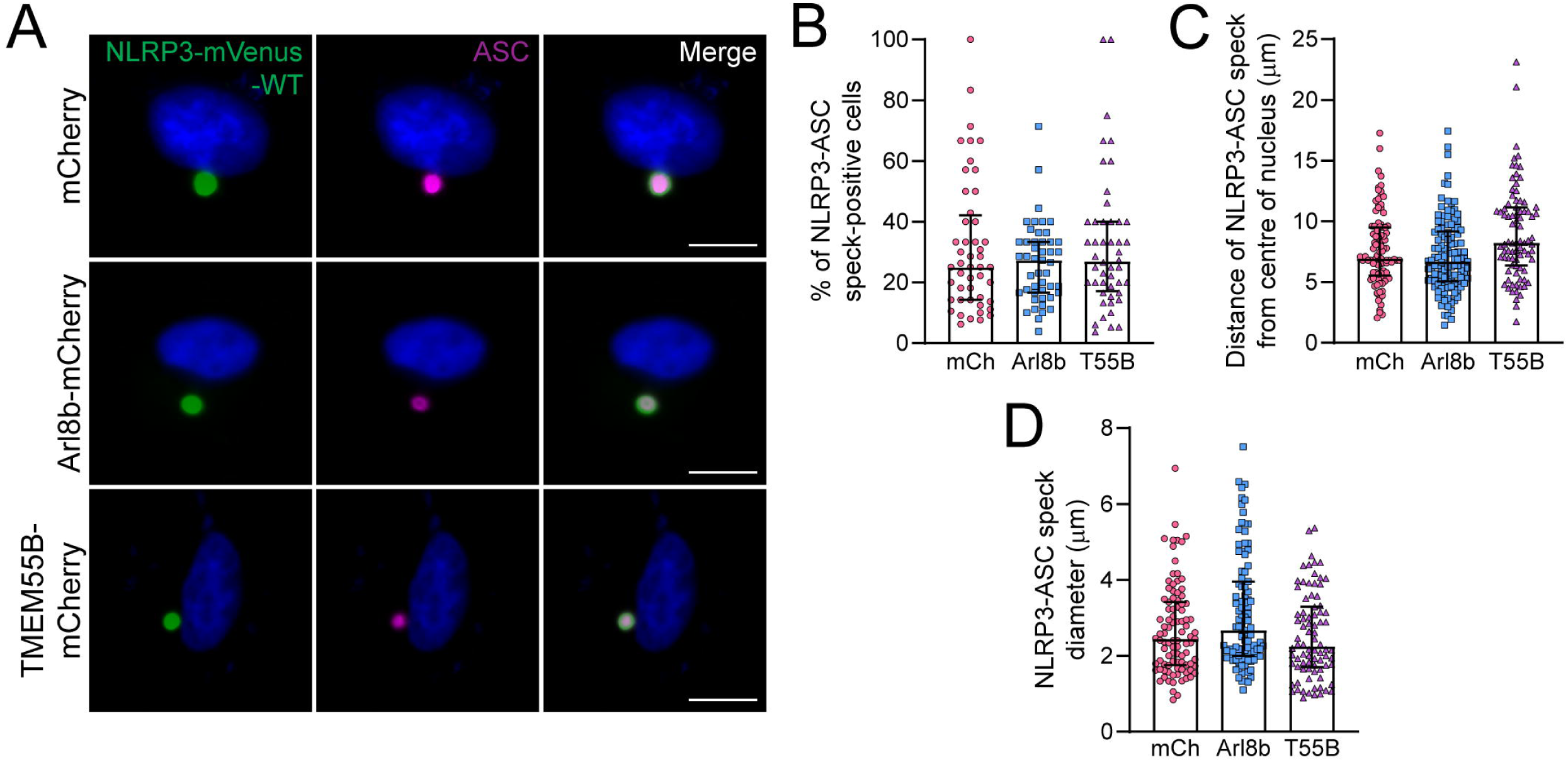
Lysosome positioning does not influence ASC speck subcellular location and size in HeLa cells. **[A]** Representative fluorescence images of all three HeLa cell lines following transfection with a combination of full-length ASC (magenta – following staining with anti-ASC antibody) and NLRP3-mVenus-WT (green). Scale bar = 10 μm. Images are representative of 48 random fields of view per cell line per treatment from four independent experiments. **[B]** Quantification of the percentage of cells positive for an NLRP3-ASC speck in all cell lines (*n* = 48 random fields of view per cell line per treatment from four independent experiments). **[C]** Quantification of the distance of the NLRP3-ASC speck from the centre of the nucleus in all cell lines. *n* = 100 (mCherry), 119 (Arl8b-mCherry), 84 (TMEM55B-mCherry) from four independent experiments. **[D]** Quantification of the diameter of the NLRP3-ASC speck in all cell lines. *n* = 95 (mCherry), 99 (Arl8b-mCherry), 82 (TMEM55B-mCherry) from four independent experiments. Data were analysed using Kruskal-Wallis test with Dunn’s multiple comparison test.

To assess effects of lysosome position on the assembly of endogenous ASC specks, Arl8b-overexpressing iBMDM cell lines were treated with nigericin in the presence of the caspase-1/4 inhibitor VX-765, included to prevent cell death (Fig 7). The ASC speck was localised further from the nucleus in Arl8b-overexpressing iBMDMs compared to control (Fig 7A and B), suggesting that lysosome positioning influences ASC speck localisation within the cell. Furthermore, the ASC speck diameter (Fig 7C) and fluorescence intensity (Fig 7D) were increased in Arl8b-overexpressing iBMDMs compared to control, suggesting that lysosome positioning also influences the size of the ASC speck in response to nigericin. We initially speculated whether the increased ASC speck size and altered position were factors contributing to the observed increased caspase-1 activation and IL-1β release in Arl8b expressing cells stimulated with nigericin (Fig 5). Thus, we treated the control and Arl8b-overexpressing iBMDMs with the additional NLRP3 activators imiquimod and ATP. Neither ATP or imiquimod caused greater caspase-1 activation in Arl8b-overexpressing cells, although IL-1β was reduced in response to ATP (Fig S3). In response to both stimuli, Arl8b-overexpressing iBMDMs formed ASC specks further away from the centre of the nucleus (Fig 7E and G, respectively), although the diameter of the ASC speck remained unchanged (Fig 7F and H). Thus, although the ASC speck was more peripherally localised upon Arl8b ovexpression and treatment with all stimuli, only nigericin resulted in a significantly increased ASC speck size. This observation suggests that increased ASC speck size could potentially account for the increased inflammasome activation observed in response to Arl8b expression and nigericin simulation. Thus, the data from iBMDM cells suggest that lysosome positioning influences the subcellular localisation and size of the ASC speck.

**Figure 7.**
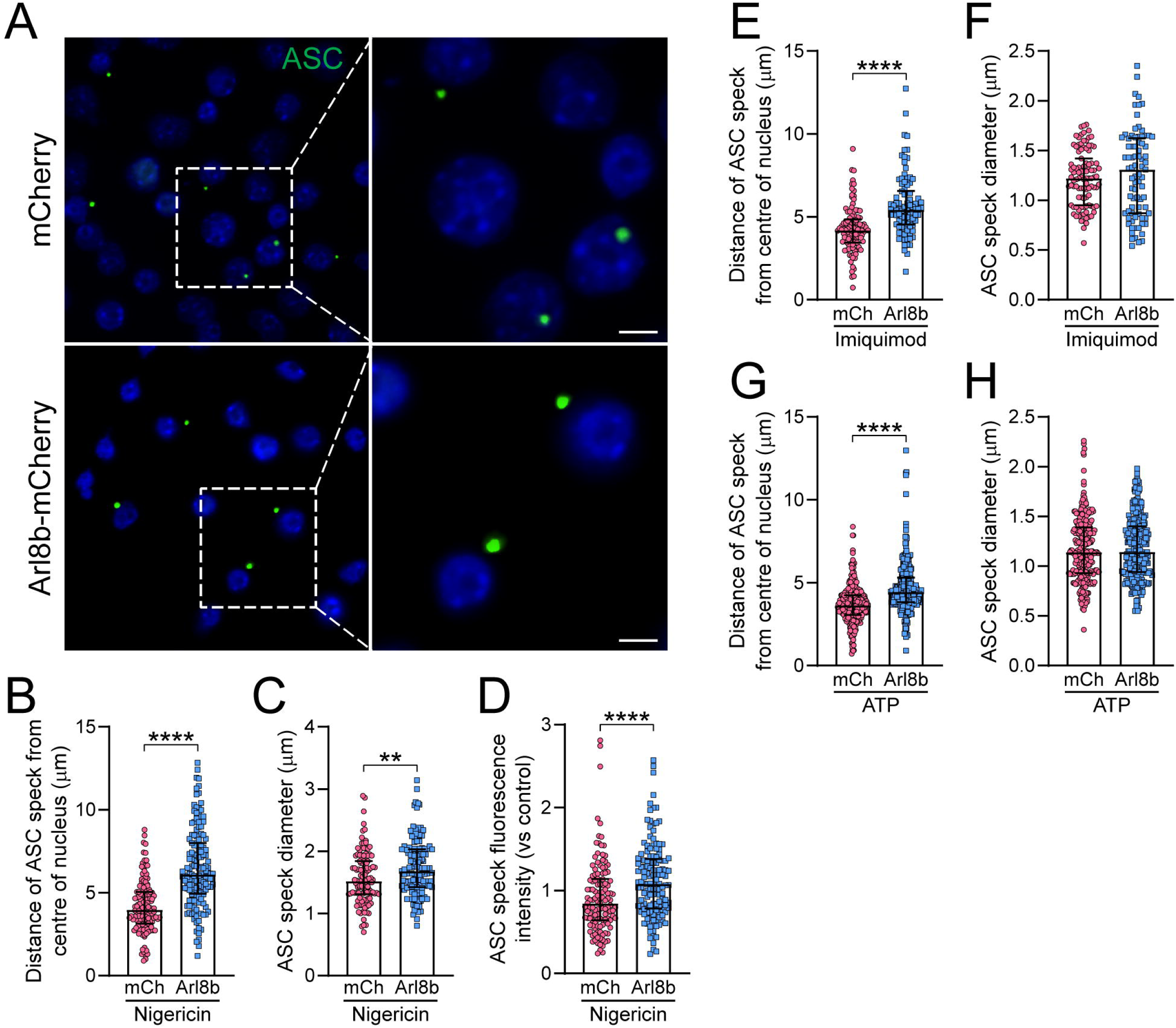
Lysosome positioning influences ASC speck subcellular location and size in iBMDMs. **[A-H]** iBMDM mCherry and iBMDM Arl8b-mCherry cells were primed with LPS (1 µg mL^−1^) in complete media for 2 h. iBMDMs were then incubated with VX765 (10 µM) for 15 min before stimulation with nigericin (10 µM) for 1 h. **[A]** Representative fluorescence images showing ASC speck distance and size (green) in each iBMDM cell line. Scale bar = 5 μm. Images are representative of 60 random fields of view per cell line per treatment from five independent experiments. **[B]** Quantification of the distance of the ASC speck from the centre of the nucleus in both iBMDM cell lines. *n* = 180 (mCherry) and 135 (Arl8b-mCherry) from five independent experiments. **[C]** Quantification of the diameter of the ASC speck in both iBMDM cell lines. *n* = 170 (mCherry) and 123 (Arl8b-mCherry) from five independent experiments. **[D]** Quantification of the ASC speck fluorescence intensity in both iBMDM cell lines. *n* = 125 (mCherry) and 135 (Arl8b-mCherry) from five independent experiments. **[E-H]** iBMDM mCherry and iBMDM Arl8b-mCherry cells were primed with LPS (1 µg mL^−1^) in complete media for 2 h. iBMDMs were then incubated with VX765 (10 µM) for 15 min before stimulation with imiquimod (150 µM) or ATP (5 mM) for 1 h. **[E]** Quantification of the distance of the ASC speck from the centre of the nucleus in both iBMDM cell lines following stimulation with imiquimod. *N* = 109 (mCherry) and 90 (Arl8b-mCherry) from five independent experiments. **[F]** Quantification of the diameter of the ASC speck in both iBMDM cell lines following stimulation with imiquimod. *n* = 99 (mCherry) and 81 (Arl8b-mCherry) from five independent experiments. **[G]** Quantification of the distance of the ASC speck from the centre of the nucleus in both iBMDM cell lines following stimulation with ATP. *n* = 294 (mCherry) and 287 (Arl8b-mCherry) from five independent experiments. **[H]** Quantification of the diameter of the ASC speck in both iBMDM cell lines following stimulation with ATP. *n* = 284 (mCherry) and 284 (Arl8b-mCherry) from five independent experiments. Data were analysed using unpaired two-tailed Mann-Whitney test. **P ≤ 0.01, ****P ≤ 0.0001.

## Discussion

The importance of lysosome positioning with respect to lysosome function is becoming increasingly recognised (Cabukusta and Neefjes, 2018, Pu et al., 2016). This is well documented in autophagy (Korolchuk et al., 2011, Willett et al., 2017) and tumour biology (Dykes et al., 2016, Glunde et al., 2003), and here we provide evidence for a role of lysosome position in the regulation of the NLRP3 inflammasome. The data presented show that inflammasomes are formed further from the centre of the nucleus in macrophages with more peripherally localised lysosomes upon nigericin activation and that, in response to nigericin, these inflammasomes are larger than those formed in control macrophages. The increase in inflammasome size could potentially account for the increased caspase-1 activation and IL-1β release observed following nigericin stimulation. The differences in IL-1β secretion in Arl8b-overexpressing cells in response to different NLRP3 stimuli are harder to reconcile, although it may be related to reduced pro-IL-1β levels seen in these cells following priming, though further research must be undertaken to investigate this observation.

We generated HeLa and iBMDM cell lines in which lysosome position was altered by the overexpression of Arl8b or TMEM55B to redistribute lysosomes to the cell periphery or perinuclear region, respectively. These manipulations were well tolerated by the cells except for overexpression of TMEM55B in iBMDMs, which were not viable (data not shown). Endosome cargo trafficking is disrupted in macrophages in response to NLRP3 inflammasome-activating stimuli (Lee et al., 2023, Zhang et al., 2023), and so it is possible that disrupted cargo trafficking could contribute to the effect of altered lysosome position upon NLRP3 inflammasome activation. Indeed, using an EGF endosome-to-lysosome trafficking assay (Liang et al., 2008, Noakes et al., 2011), defects in cargo trafficking were observed in HeLa cells with manipulated lysosome localisation. As lysosome position is known to play a role in lysosomal homeostasis (Johnson et al., 2016, Pu et al., 2016), the effects of lysosome position on lysosomal pH and health were investigated. Similar to previous findings (Johnson et al., 2016), peripheral lysosomes had higher luminal pH, whereas perinuclear lysosomes had slightly lower luminal pH. Maintenance of an acidic luminal pH is essential for lysosome function, and dysregulation of lysosomal acidification can lead to impaired cellular homestasis, such as disruption to endocytosis and autophagy, as well as macromolecule biogenesis and transport (Lawrence and Zoncu, 2019), all of which can contribute to conditions such as proteinopathic neurodegenerative diseases, metabolic disorders, and immunological diseases (Zeng et al., 2020). In addition, dysregulated lysosomal acidification also affects other organelles such as mitochondria, which can lead to increased reactive oxygen species production and pro-inflammatory cytokine release, contributing to the pathogenesis of inflammatory disease and cancer (Deus et al., 2020, Yambire et al., 2019, Zeng et al., 2020).

Manipulation of lysosome position in HeLa cells and macrophages caused lysosomal damage at baseline, as indicated by increased number of cells positive for galectin-3 puncta. As lysosomal damage is known to contribute to NLRP3 inflammasome activation (Hornung et al., 2008, Lima et al., 2013, Munoz-Planillo et al., 2013), altered lysosome positioning may contribute to inflammasome activation by increasing or sensitising lysosomal damage. Other evidence suggesting lysosome positioning could be important for NLRP3 inflammasome activation is the reported role of LAMTOR, which interacts with BORC to restrain lysosomes to the perinuclear region, thus negatively regulating Arl8b-dependent peripheral positioning (Filipek et al., 2017). LAMTOR also interacts directly with NLRP3, an interaction suggested to be required for NLRP3 inflammasome activation (Tsujimoto et al., 2023).

The NLRP3 inflammasome is activated at sites of tissue injury, infection, or disease pathology, where inflammatory stressors disrupt cell physiology and homeostasis (Liston and Masters, 2017, Seoane et al., 2020), conditions that likely contribute to redistribution of lysosomes within the cell. An acidic extracellular environment, such as the acidic tumour microenvironment, alters both lysosome function and position, driving lysosomes to the cell periphery and promoting tumorigenesis (Glunde et al., 2003). Extracellular acidosis also characterises sites of inflammation, and is known to influence immune cell function (Lardner, 2001). Therefore, tissue acidosis at sites of inflammation may influence inflammasome signalling through repositioning of lysosomes. Thus, there is a need to understand how subcellular organisation and organelle position influence inflammatory signalling in response to different stimuli.

In summary, the data presented here highlight a role for lysosome positioning in NLRP3 inflamasome activation. Why repositioning lysosomes to the cell periphery increased NLRP3 inflammasome activation in response to nigericin requires further investigation, but may be related to the formation of the ASC speck further from the centre of the nucleus and the size of the ASC speck. The new insight delivered here opens new paths of investigation that will continue to further our understanding of the mechanisms regulating NLRP3 inflammasome activation, and open new avenues for therapeutic intervention in diseases where lysosome position is modified.

## Acknowledgments

This work was funded by a Medical Research Council (MRC) doctoral training programme (MR/N013751/1) to B.J.M., an MRC grant to D.B. and M.L. (MR/T016515/1), and an MRC grant (MR/T016043/1) and BBSRC grant (BB/Y004876/1) to G.LC. The Bioimaging Facility microscopes used in this study were purchased with grants from BBSRC, Wellcome and the University of Manchester Strategic Fund. Special thanks goes to Peter March and Steven Marsden for their help with the microscopy.

## Author contributions

Conceptualization D.B., M.L., G.LC.; Methodology H.B., A.A.,; Formal Analysis B.J.M., C.H., D.B.; Investigation B.J.M., C.H., J.P.G., P.I.S., T.D., A.O.; Resources H.B., A.A.; Data Curation B.J.M., C.H.; Writing – Original Draft Preparation B.J.M., D.B., M.L., C.H.; Writing – Review & Editing Preparation all authors; Visualization B.J.M., C.H.; Supervision D.B., M.L., G.LC., C.H., P.I.S., J.P.G.; Project Administration D.B., M.L., G.LC.; Funding Acquisition D.B., M.L., G.LC.

## Declaration of interests

The authors have no competing interests.

## Supplemental Information

Document S1. Figures S1-S3 Document S2. Supplementary methods

## STAR Methods

### Key resources table

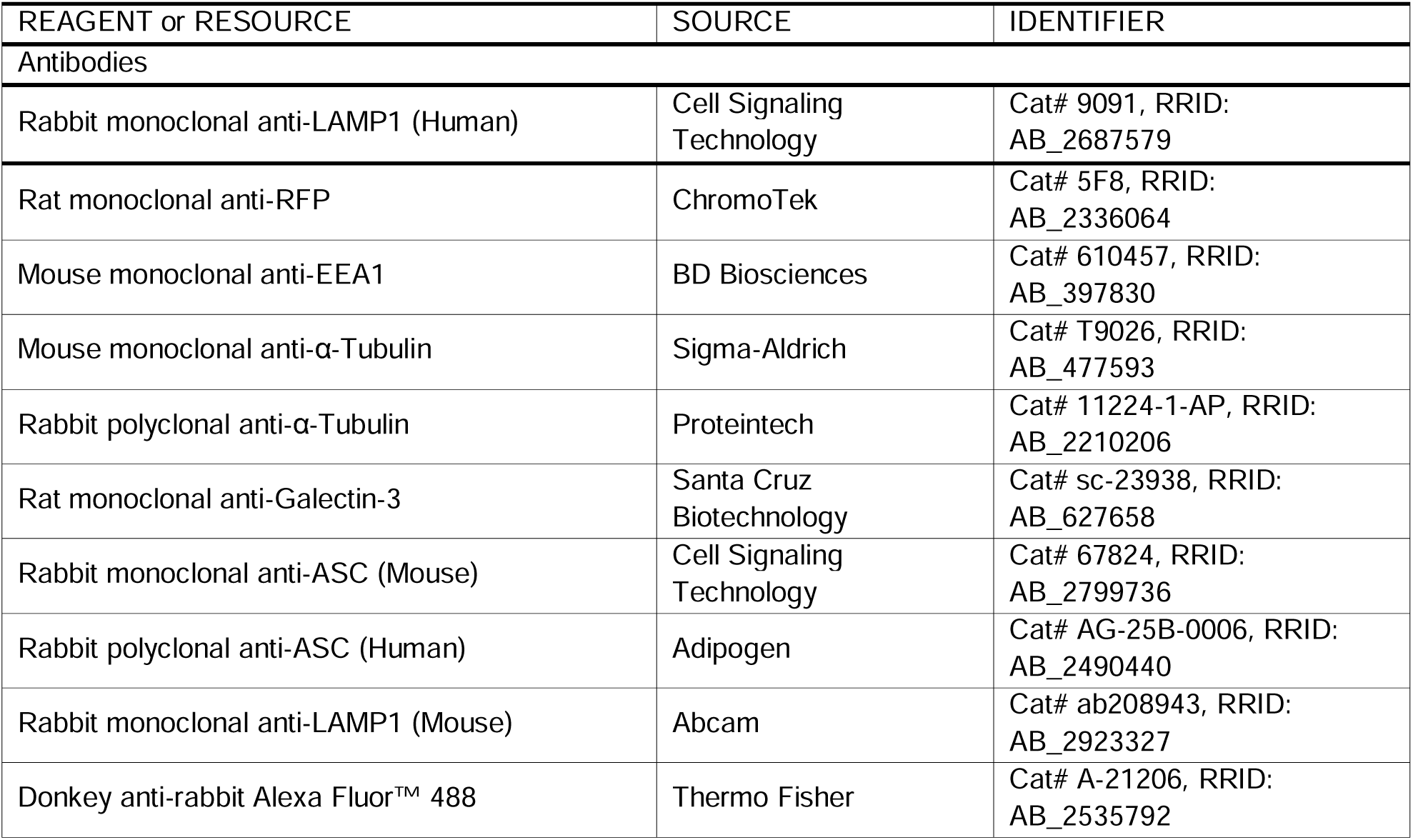
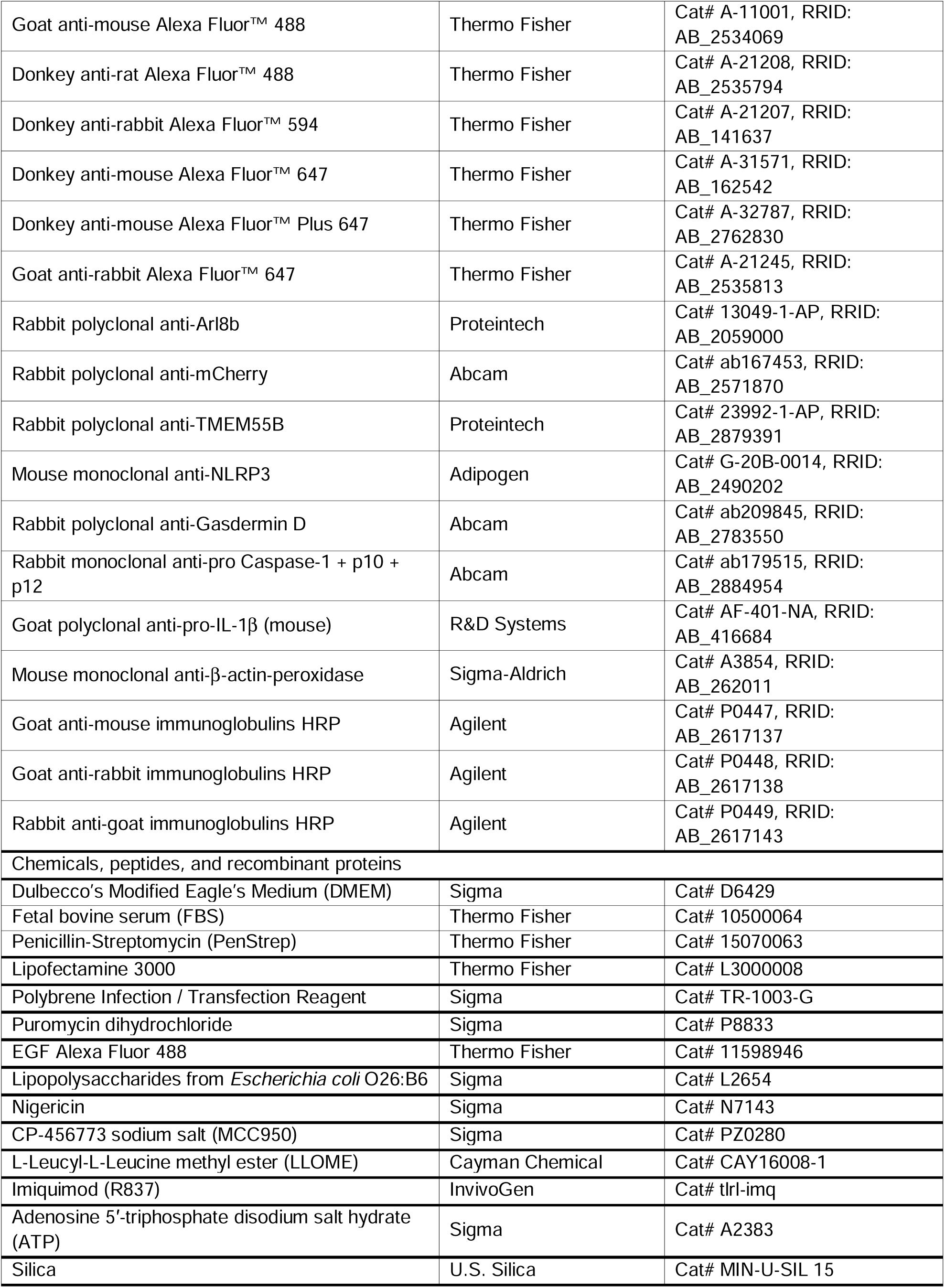
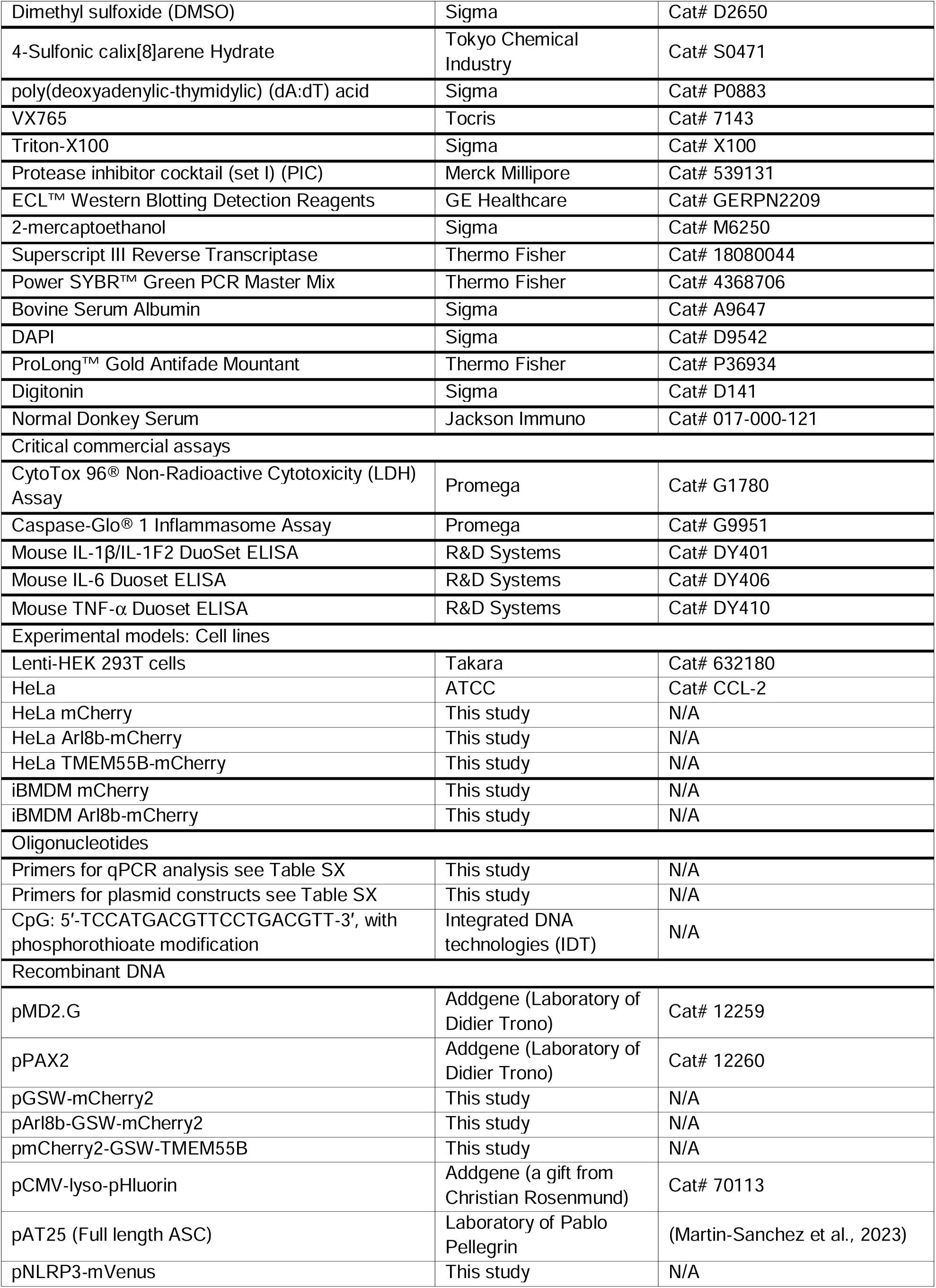
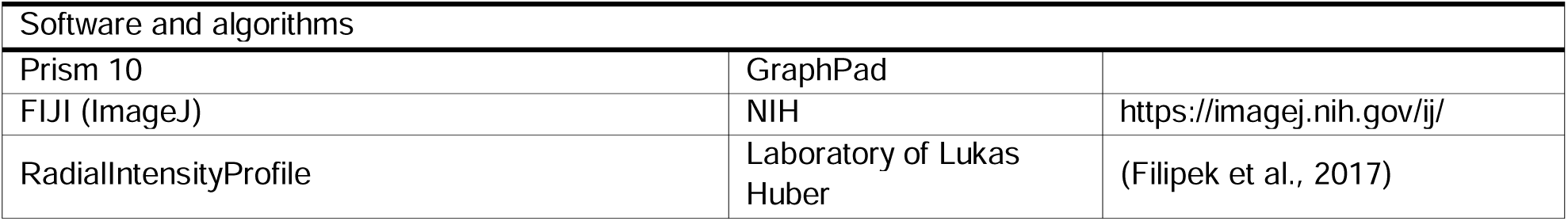

### Experimental model and study participant details

#### Cell culture

Lenti-HEK 293T cells were purchased from Takara. HeLa cells were purchased from ATCC. HeLa mCherry, HeLa Arl8b-mCherry, HeLa TMEM55B-mCherry, iBMDM mCherry and iBMDM Arl8b-mCherry were generated as part of this study. All cells were cultured in complete media, which consists of Dulbecco’s modified Eagle’s medium (DMEM) containing 10% (v/v) fetal bovine serum (FBS) and 1% (v/v) Penicillin-Streptomycin (PenStrep). Transduced cell lines were cultured in complete media containing 1.5 µg mL^−1^ puromycin. All cell lines were maintained at 37°C and 5% CO_2_. Before experiments, cells were seeded at the density stated in each experiment and left to adhere overnight at 37°C and 5% CO_2_.

## Method details

### Generation of plasmids and cell lines

All vectors were created by the Genome Editing Unit at The University of Manchester (see supplementary methods). Hifi assembly fragments were amplified with HF KOD polymerase. Lentiviral vectors were generated using NEB Gibson HiFi assembly kit (New England BioLabs) according to manufacturer’s instructions, in a 3^rd^ generation lentiviral backbone. Plasmid maps are provided in the supplementary methods for each vector.

Lenti-HEK 293T cells were plated at 0.5 x 10^6^ cells per well in a 1% (v/v) poly-L-lysine-coated 6-well plate to promote adherence. After 24 h, Lenti-HEK 293T cells were transfected with 1.2 μg pMD2.G, 0.4 μg pPAX2 and 1.5 μg of either GSW-mCherry2, Arl8b-GSW-mCherry2, mCherry2-GSW-TMEM55B or PBS as a mock-transduced control using Lipofectamine 3000 and performed following the manufacturer’s protocol. Following 24 h of transfection, media was replaced with fresh DMEM, and the cells were incubated for a further 48 h. Viral suspensions were then collected and filtered using a 0.45 μm filter to remove any detached cells. The filtered viral suspension was then used to transduce HeLa cells plated at 0.1 x 10^6^ cells mL^−1^ in a 12-well plate. To do so, HeLa cells were incubated with viral suspension containing 0.8 µg mL^−1^ polybrene for 6 h before media was replaced with fresh DMEM. HeLa cells were incubated at 37°C for 4 days before media was replaced with DMEM containing 1.5 µg mL^−1^ puromycin to begin selection. GSW-mCherry2, Arl8b-GSW-mCherry2 and mCherry2-GSW-TMEM55B transduced cells (named HeLa mCherry, HeLa Arl8b-mCherry and HeLa TMEM55B-mCherry, respectively, from herein) and mock-transduced control were maintained in puromycin media until all non-transduced cells were visibly dead under light microscope. Visual inspection confirmed mCherry expression in cell lines confirmed by western blotting.

iBMDM overexpressing cell lines were generated using the same method as above. Filtered GSW-mCherry2, Arl8b-GSW-mCherry2, mCherry2-GSW-TMEM55B and mock viral suspensions containing 0.8 µg mL^−1^ polybrene were used to transduce iBMDM cells plated at 0.1 x 10^6^ cells mL^−1^ in a 12-well plate. mCherry2-GSW-TMEM55B overexpression in iBMDMs caused cell death to the extent that the cell line was no longer viable, suggesting that perinuclear lysosome positioning in macrophage exerts a toxic phenotype. Therefore, we were unable to obtain this cell model. GSW-mCherry2 and Arl8b-GSW-mCherry2 transduced cells (named iBMDM mCherry and iBMDM Arl8b-mCherry from herein) were taken forward for further experiments.

### Trafficking experiments

The EGF trafficking experiment was conducted as described previously (Lee et al., 2023). HeLa cells were seeded at 0.1 x 10^6^ cells mL^−1^ on coverslips. Cells were washed three times with 1X PBS before incubation with 0.4 µg mL^−1^ EGF Alexa Fluor 488 in serum-free DMEM supplemented with 2% (w/v) BSA on ice for 1 h. Cells were then washed twice with cold 1X PBS before incubation with DMEM at 37°C for 1-2 h.

### Lysosomal health and function

To assess lysosomal damage, HeLa cells were seeded at 0.1 x 10^6^ cells mL^−1^ and iBMDM cells were seeded at 0.375 x 10^6^ cells mL^−1^ on coverslips. iBMDMs were then primed with 1 µg mL^−1^ LPS in complete DMEM for 2 h. Following this, the cell media was replaced with serum-free media and cells were treated with vehicle (ethanol; 0.5% v/v) or LLOMe (1 mM) for 30 min, followed by processing for immunocytochemistry and co-staining with an anti-galectin-3 antibody. To assess lysosomal pH, HeLa cells were seeded at 0.1 x 10^6^ cells mL^−1^ on coverslips. Cells were transfected with 0.5 μg pCMV-lyso-pHluorin using Lipofectamine 3000 for 16 h, followed by co-staining with an anti-LAMP-1 antibody.

### Inflammasome activation experiments

iBMDM cells were seeded at a density of 0.75 x 10^6^ cells mL^−1^ in complete DMEM and left to adhere overnight. For NLRP3 experiments, cells were primed with LPS (1 µg mL^−1^) in complete media for 2 h. To assess the effects of lysosome positioning on NLRP3 inflammasome activation, the cell media was replaced with serum-free media containing vehicle (PBS) or MCC950 (10 µM), a NLRP3-specific inhibitor, for 15 min. Cells were then stimulated with a range of NLRP3-activating stimuli. For nigericin experiments, cells were stimulated with vehicle (ethanol; 0.5% v/v) or nigericin (10 µM) for 1 h. For imiquimod experiments, cells were stimulated with vehicle (H_2_O; 1% v/v) or imiquimod (150 µM) for 2 h. For ATP experiments, cells were stimulated with vehicle (H_2_O; 1% v/v) or ATP (5 mM) for 1 h. For silica experiments, cells were incubated with vehicle (PBS) or MCC950 (10 µM) concurrently with vehicle (PBS; 0.5% v/v) or silica (300 μg mL^−1^) for 4 h. To assess the effects of lysosome positioning on AIM2 inflammasome activation in iBMDMs, the media was replaced with serum-free media containing vehicle (DMSO; 0.5% v/v) or 4-Sulfonic calix[8]arene (10 µM) for 15 min following cell priming. Cells were then activated by lipofectamine 3000-mediated transfection of mock (PBS) or poly(dA:dT) (1 µg mL^−1^) for 4 h. To assess ASC speck number, iBMDMs were seeded at 0.5 × 10^6^ cells mL^−1^ on coverslips. iBMDMs were primed with 1 µg mL^−1^ LPS in complete DMEM for 2 h. iBMDMs were then incubated with VX765 (10 µM) with or without MCC950 (10 µM) in serum-free DMEM for 15 min. Cells were then treated with vehicle (ethanol; 0.5% v/v) or nigericin (10 µM) for 1 h before processing for immunocytochemistry and staining with an anti-ASC antibody.

### NLRP3-mVenus-WT and ASC transfection experiment

To assess the effects of lysosome positioning on ASC speck formation, HeLa cells were seeded at 0.1 x 10^6^ cells mL^−1^ on coverslips. Cells were transfected with either 0.2 μg full-length ASC, 0.1 μg NLRP3-mVenus-WT, or a combination of both using Lipofectamine 3000 for 16 h, followed by staining with an anti-ASC antibody. For the iBMDM experiments, cells were seeded at 0.375 × 10^6^ cells mL^−1^ on coverslips. iBMDMs were primed with 1 µg mL^−1^ LPS in complete DMEM for 2 h. iBMDMs were then incubated with VX765 (10 µM) in serum-free DMEM for 15 min. Cells were then treated with nigericin (10 µM), ATP (5 mM) or imiquimod (150 µM) for 1 h before immunocytochemistry with an anti-ASC antibody.

### Cell death assay (LDH)

Supernatants were measured for lactate dehydrogenase (LDH) release as a determinant of cell death using the CytoTox 96 nonradioactive cytotoxicity assay, according to the manufacturer’s instructions.

### Caspase-1 activity

Supernatants were measured for caspase-1 activity using Caspase-Glo® 1 Inflammasome Assay. Briefly, supernatants were combined with Z-WEHD aminoluciferin substrate and incubated for 1 h at room temperature (RT) before measuring luminescence using a Biotek Synergy HT Microplate Reader.

### ELISA

For inflammasome activation, supernatants were analysed for IL-1β. For the priming and TLR signaling experiments, supernatants were measured for IL-6 and TNF-α. All ELISAs were carried out according to manufacturer’s instructions.

### Western blot

Cells were lysed with lysis buffer (50 mM Tris-HCl, 150 mM NaCl, 1% (v/v) Triton x-100), containing protease inhibitor cocktail on ice for 15 min. Cell lysates were clarified by centrifugation for 15 min at 12,000 *x g* at 4°C before the supernatant was removed and combined with 5X Laemmli buffer. Equal volumes of cell lysate samples were run on SDS-polyacrylamine gels and transferred at 25 V onto PVDF membranes using a Trans-Blot^®^ Turbo Transfer™ System (Bio-Rad). Membranes were blocked in 5% (w/v) BSA in PBS, 0.1% Tween-20 (PBST) for 1 h at RT. Membranes were then incubated at 4°C overnight with primary antibodies in 1% (w/v) BSA in PBST. Membranes were washed with PBST and incubated with secondary antibodies for 1 h at RT. Following incubation with Amersham ECL Western Blotting Detection Reagent, proteins were visualized using G:BOX (Syngene) and Genesys software. β-actin was used as a loading control.

### qPCR

iBMDMs were lysed with RNA lysis buffer containing 1% (v/v) 2-mercaptoethanol on ice and RNA was extracted using the PureLink™ RNA Mini Kit. After RNA extraction, cDNA was synthesized using Superscript III Reverse Transcriptase (according to the manufacturer’s instructions). Primers were designed using Primer-BLAST and ordered from Invitrogen. Primer sequences used are detailed in the supplementary methods. Samples were run in triplicate using *Power* SYBR™ Green PCR Master Mix and 200 nM of each primer on a 7900HT Fast Real-Time PCR System (Applied Biosystems). Relative expression levels were calculated using the 2^−ΔΔCt^ method normalising to the expression of the housekeeping gene *Hmbs*.

### Immunocytochemistry

For lysosomal and endosomal positioning characterisation experiments, HeLa cells were seeded in a 24-well plate at 0.05 x 10^6^ cells mL^−1^ on coverslips and left to adhere overnight at 37°C and 5% CO_2_. For initial characterisation experiments, HeLa cells were stained using anti-LAMP1 and anti-RFP antibodies. For all subsequent characterisation experiments, HeLa cells were stained using an anti-α-tubulin antibody. For LAMP1 staining, cells were washed once with 1X PBS and fixed in ice-cold 100% methanol for 6 min on ice. Cells were washed once and blocked with 5% BSA (w/v) in PBS for 1 h at RT. For EEA1 staining, cells were fixed with 4% (w/v) PFA for 15 min at RT. Cells were washed once with 1X PBS and then permeabilised with 0.2% (v/v) Triton-X-100 in PBS for 5 min.

For all immunocytochemistry experiments, cells were then washed once and blocked with 5% (w/v) BSA in PBS for 1 h at RT. Cells were then incubated with the desired primary antibody at a pre-determined concentration in 5% (w/v) BSA in PBS for 1 h at RT. Cells were washed twice with 1X PBS before incubation with the desired secondary antibody in 5% (w/v) BSA in PBS for 1 h at RT. Cells were washed twice in 1X PBS and incubated with DAPI (0.5 µg mL^−1^) for 10 min at RT before final washing in dH_2_O and mounting using ProLong™ Gold Antifade Mountant prior to imaging using widefield microscopy.

For lysosome positioning experiments in iBMDMs, cells were seeded in a 24-well plate at 0.2 x 10^6^ cells mL^−1^ on coverslips and left to adhere overnight at 37°C and 5% CO_2_. Following this, iBMDMs were primed with LPS (1 µg mL^−1^) in complete DMEM for 2 h. Cells were washed twice with 1X PBS containing Ca^2+^ and Mg^2+^ before fixing with 4% (w/v) PFA for 15 min at RT. Cells were then washed twice with 1X PBS containing 50 mM NH_4_Cl before permeabilisation with 20 µM digitonin in PIPES buffer (20 mM PIPES, 137 mM NaCl, 2.7 mM KCl, pH 6.8) for 10 min at RT. Cells were then blocked with 5% (v/v) donkey serum in 1X PBS containing 50 mM NH_4_Cl for 1 h at RT. Cells were incubated with anti-LAMP1 and anti-α-tubulin primary antibodies in 5% (v/v) donkey serum in 1X PBS containing 50 mM NH_4_Cl overnight at 4°C. Following this, cells were washed thrice with PIPES buffer before incubation with the desired secondary antibodies in 5% (v/v) donkey serum in 1X PBS containing 50 mM NH_4_Cl for 1 h at RT. Cells were then washed once with PIPES buffer and incubated with DAPI (0.5 µg mL^−1^) for 10 min at RT. Cells were washed once with PIPES buffer and then fixed with 2% (w/v) PFA for 5 min at RT. Cells were then washed thrice with 1X PBS containing 50 mM NH_4_Cl, washed thrice with dH_2_O and then left to dry for 2 h before mounting using ProLong™ Gold Antifade Mountant prior to imaging using widefield microscopy.

### Immunofluorescence microscopy

Widefield immunofluorescence microscopy was used throughout this study. Images were obtained on a Zeiss Axioimager.D2 Upright microscope using a 63x Plan Apochromat objective, except for iBMDM ASC speck number experiments, which used a 20x EC Plan-neofluar. Images were captured using a Coolsnap HQ2 camera (Photometrics) through Micromanager software (v1.4.23). Specific band pass filter sets for DAPI, FITC, Texas Red and Cy5 were used to prevent bleed through from one channel to the next.

## Quantification and statistical analysis

### Image processing analysis

For lysosomal and endosomal positioning experiments, images were taken of at least 10 independent fields of view within each independent experimental repeat. For EGF trafficking, lysosomal damage and pHluorin experiments, images were taken of 6 independent fields of view per cell line per treatment within each experimental repeat. For the HeLa NLRP3-mVenus transfection, images were taken of 12 independent fields of view per cell line per treatment within each experimental repeat. To assess ASC speck distance and size in HeLa cells and iBMDMs, images were taken of 12 independent fields of view per cell line per treatment within each experimental repeat. For ASC speck number experiments in iBMDMs, images were taken of 5 independent fields of view per cell line per treatment within each experimental repeat.

### Image processing analysis

Image processing and analysis was performed using Fiji (ImageJ). For quantification of lysosomal and endosomal positioning, a RadialIntensityProfile macro was used to analyze the radial distribution of fluorescence signal in relation to the cell nucleus (Barral et al., 2022, Filipek et al., 2017). For HeLa cells, fluorescence intensity was determined at 2 µm increments from the center of the nucleus to 24 µm away from the nucleus. For iBMDM cells, fluorescence intensity was determined at 1.3 µm increments from the center of the nucleus to 20.6 µm away from the nucleus. Any fluorescence intensity less than or equal to 8 µm (6.5 µm for iBMDMs) from the center of the nucleus was deemed “perinuclear”, whereas fluorescence intensity more than 8 µm (6.5 µm for iBMDMs) from the center of the nucleus was deemed “peripheral”. For lysosome positioning analysis in HeLa cells, 96-110 cells per cell line were analysed from 3 independent repeats. For lysosome positioning analysis in iBMDM cells, 159 cells per cell line were analysed from 4 independent repeats. For endosome positioning analysis, 90 cells per cell line were analysed from 3 independent repeats. For lysosome positioning in NLRP3-mVenus-WT experiments, 64 cells per cell line per condition were analysed from 4 independent experiments. For lysosomal damage experiments, the percentage of cells in the field of view positive for galectin-3 puncta was calculated. For lysosomal pH experiments, the total pHluorin fluorescence intensity of 56-64 cells per cell line per treatment were analysed from 4 independent experiments. For EGF trafficking assay (Lee et al., 2023), background was calculated within each image by taking independent measurements from areas where cells were absent. The average background readout was subsequently subtracted from the respective mean fluorescence to obtain a final fluorescence value. For ASC speck distance, a measurement was taken from the center of the nucleus to the center of the ASC speck using the ImageJ measuring tool. For ASC speck size, the diameter of the ASC speck was measured using the ImageJ measuring tool. For ASC speck number analysis, the percentage of cells in the field of view positive for an ASC speck was calculated. For ASC speck fluorescence intensity, background was calculated for each image and was subsequently subtracted from the respective mean fluorescence to obtain a final fluorescence value.

### Statistical analysis

All imaging data are presented as median ± interquartile range, with each individual data point shown as representation of a cell or a field of view analysed from each independent experiment. The remaining data are presented as the mean ± standard error of the mean (SEM) together with individual data points where possible. Data were assessed for normal distribution using the Shapiro-Wilk normality test. Parametric data were analysed using unmatched one-way or two-way analysis of variance (ANOVA) with Holm-Sidak’s post hoc test. For nonparametric data, unpaired two-tailed Mann-Whitney test or Kruskal-Wallis test followed by Dunn’s multiple comparison test was used. Analyses were performed using GraphPad Prism (v10). Statistical significance was accepted at \**p* < 0.05. Blots are representative of two to four independent experiments.

## Notes

### Competing Interest Statement

The authors have declared no competing interest.

