## Supplementary Figures 1-3 for "Lysosome positioning affects NLRP3 inflammasome activation"

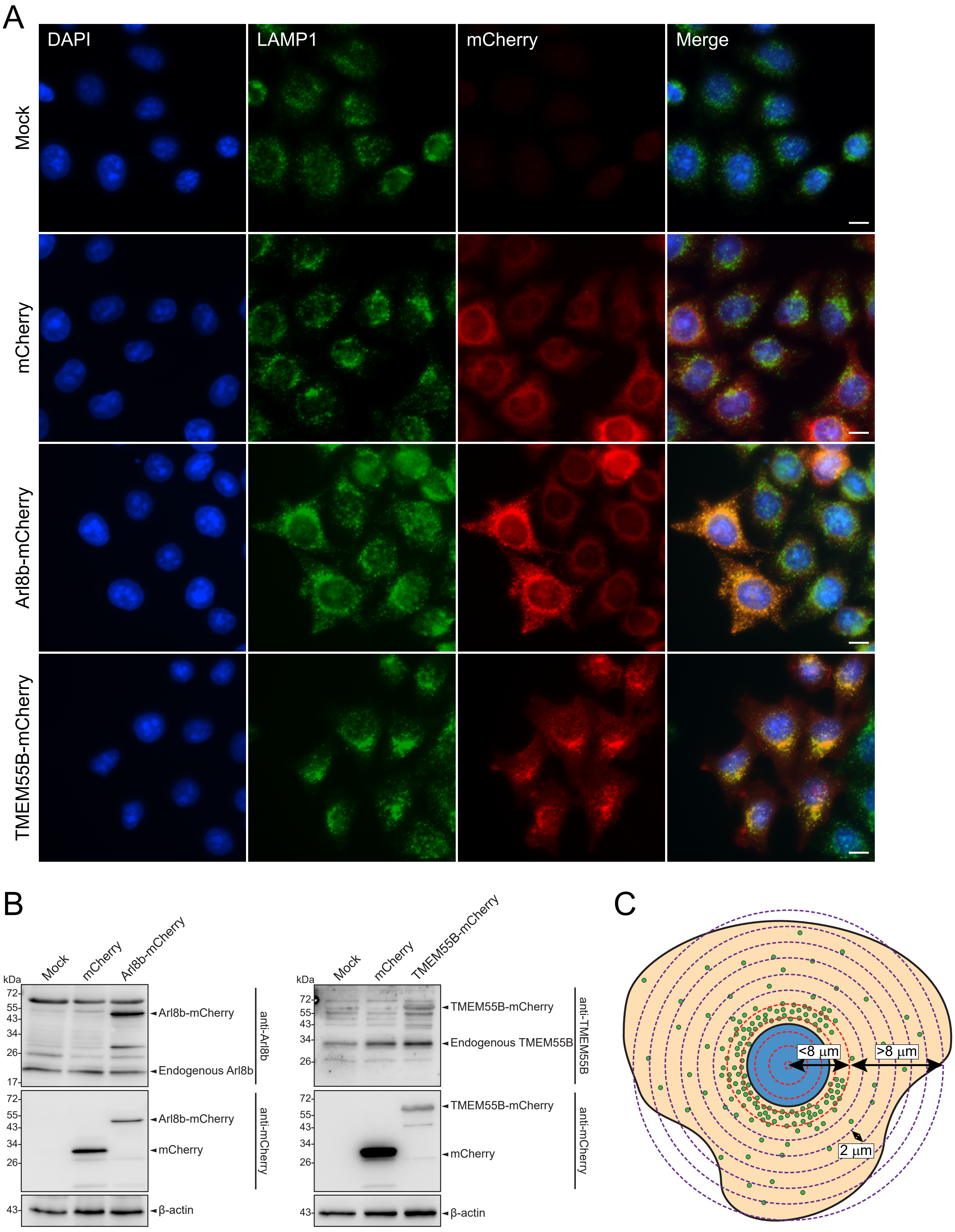
A
DAPI
LAMP1
mCherry
Merge
Mock
mCherry
Arl8b-mCherry
TMEM55B-mCherry
B
Mock
mCherry
Arl8b-mCherry
kDa
72-
55-
43-
34-
26-
17-
Arl8b-mCherry
Endogenous Arl8b
anti-Arl8b
Arl8b-mCherry
mCherry
anti-mCherry
β-actin
Mock
mCherry
TMEM55B-mCherry
TMEM55B-mCherry
Endogenous TMEM55B
anti-TMEM55B
TMEM55B-mCherry
mCherry
anti-mCherry
β-actin
C
<8 µm
>8 µm
2 µm

**Supp. Figure 1. Characterisation of HeLa mock, HeLa mCherry, HeLa Arl8b-mCherry and HeLa TMEM55B-mCherry cell lines. [A]** Representative images for HeLa mock-transduced cells, alongside HeLa mCherry, HeLa Arl8b-mCherry and HeLa TMEM55B-mCherry cells, stained for the lysosomal marker LAMP1 (green) and mCherry stained with anti-RFP (red). mCherry signal was adjusted equally between mock-transduced and mCherry alone, and differentially adjusted for Arl8b-mCherry and TMEM55B-mCherry to avoid saturation and to highlight the distribution of mCherry signal. Scale bar = 10 μm. Images are representative of 42 random fields of view per cell line from four independent experiments. **[B]** Western blots of cell lysates from mock-transduced, HeLa mCherry, HeLa Arl8b-mCherry and HeLa TMEM55B-mCherry, showing overexpression of proteins. Blots are representative of two independent experiments. **[C]** Schematic showing the ImageJ RadialIntensityProfile plugin used for quantification of the position of fluorescence signal. Fluorescence intensity measurements were taken at 2 μm increments in concentric rings from the centre of the nucleus. Fluorescence signal ≤8 μm from the centre of the nucleus was deemed perinuclear. Fluorescence signal >8 μm from the centre of the nucleus was deemed peripheral.

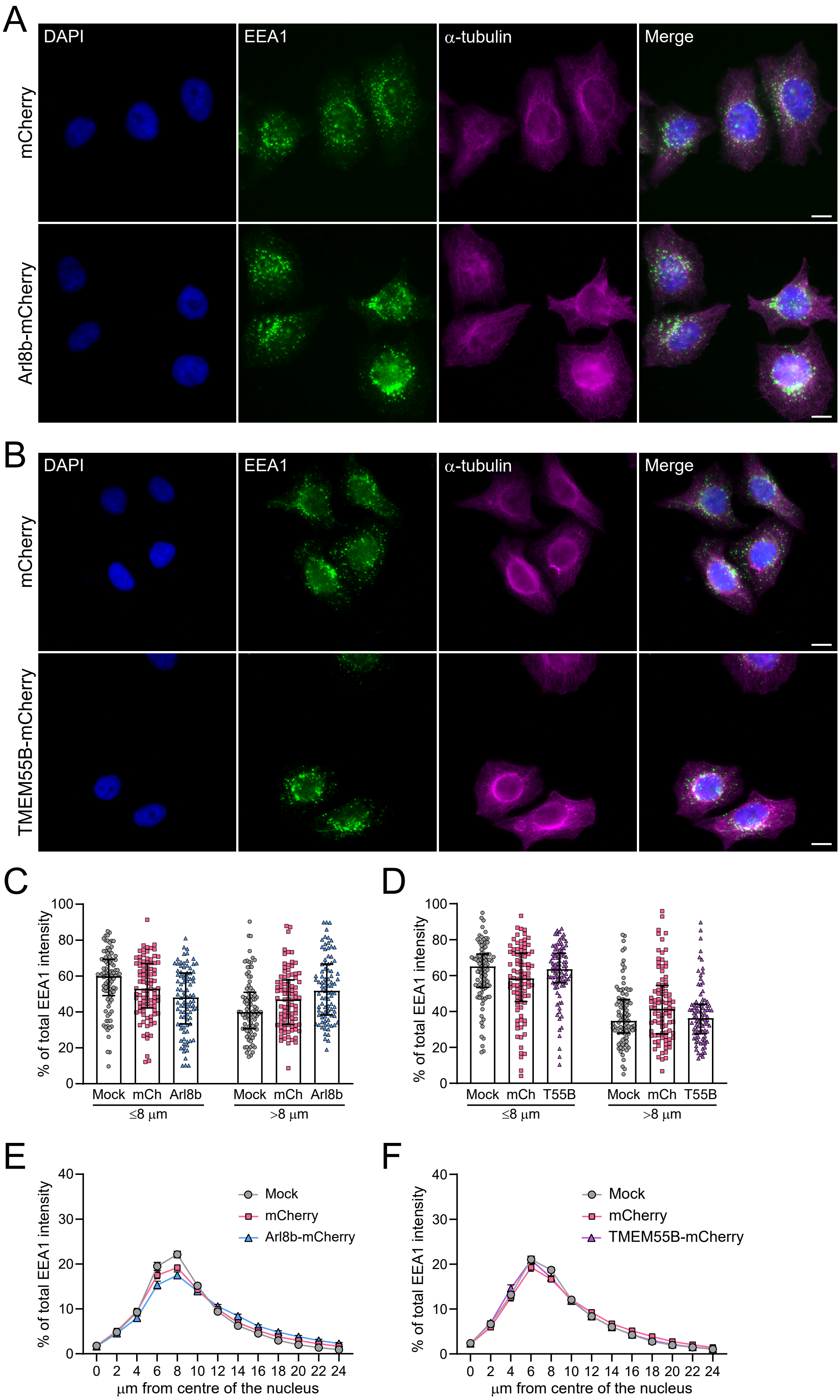

A
DAPI
EEA1
α-tubulin
Merge
mCherry
Arl8b-mCherry
B
DAPI
EEA1
α-tubulin
Merge
mCherry
TMEM55B-mCherry
C
% of total EEA1 intensity
Mock mCh Arl8b
≤8 µm
Mock mCh Arl8b
>8 µm
D
% of total EEA1 intensity
Mock mCh T55B
≤8 µm
Mock mCh T55B
>8 µm
E
% of total EEA1 intensity
Mock
mCherry
Arl8b-mCherry
µm from centre of the nucleus
F
% of total EEA1 intensity
Mock
mCherry
TMEM55B-mCherry
µm from centre of the nucleus

**Supp. Figure 2. Effect of Arl8b-mCherry and TMEM55B-mCherry overexpression in HeLa cells on early endosome positioning. [A and B]** Representative images for HeLa mCherry, alongside HeLa Arl8b-mCherry **[A]** and HeLa TMEM55B-mCherry **[B]**, stained for the early endosome marker EEA1 (green) and the microtubule marker $\alpha$-tubulin (magenta). Scale bar = 10 $\mu$m. Images are representative of 36 random fields of view per cell line from three independent experiments. **[C and D]** Quantification of the early endosome marker EEA1 fluorescence intensity at ≤ 8 $\mu$m and > 8 $\mu$m from the centre of the nucleus in HeLa Arl8b-mCherry **[C]** and HeLa TMEM55B-mCherry cells **[D]** (n = 90 individual cells per cell line from three independent repeats). **[E and F]** Quantification of the percentage total EEA1 fluorescence intensity at each 2 $\mu$m interval in HeLa Arl8b-mCherry **[E]** and HeLa TMEM55B-mCherry **[F]** cell lines, showing early endosome distribution within the cell (n = 90 individual cells per cell line from three independent repeats). Data were analysed using Kruskal-Wallis test with Dunn's multiple comparison test.

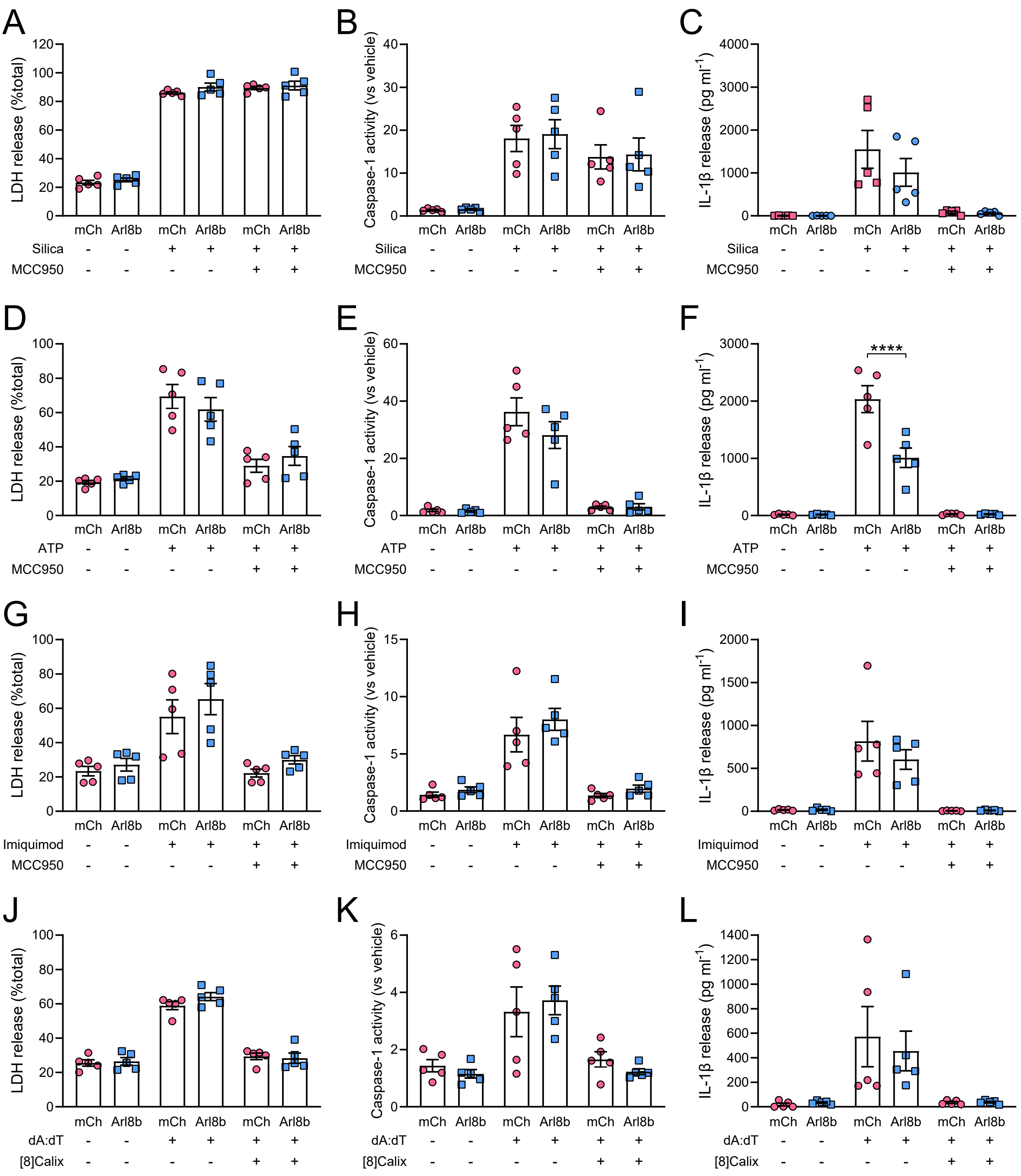
A
LDH release (%total)
0
20
40
60
80
100
120
mCh Arl8b mCh Arl8b mCh Arl8b
Silica - - + + + +
MCC950 - - - - + +
B
Caspase-1 activity (vs vehicle)
0
10
20
30
40
mCh Arl8b mCh Arl8b mCh Arl8b
Silica - - + + + +
MCC950 - - - - + +
C
IL-1β release (pg ml-1)
0
1000
2000
3000
4000
mCh Arl8b mCh Arl8b mCh Arl8b
Silica - - + + + +
MCC950 - - - - + +
D
LDH release (%total)
0
20
40
60
80
100
mCh Arl8b mCh Arl8b mCh Arl8b
ATP - - + + + +
MCC950 - - - - + +
E
Caspase-1 activity (vs vehicle)
0
20
40
60
mCh Arl8b mCh Arl8b mCh Arl8b
ATP - - + + + +
MCC950 - - - - + +
F
IL-1β release (pg ml-1)
0
1000
2000
3000
****
mCh Arl8b mCh Arl8b mCh Arl8b
ATP - - + + + +
MCC950 - - - - + +
G
LDH release (%total)
0
20
40
60
80
100
mCh Arl8b mCh Arl8b mCh Arl8b
Imiquimod - - + + + +
MCC950 - - - - + +
H
Caspase-1 activity (vs vehicle)
0
5
10
15
mCh Arl8b mCh Arl8b mCh Arl8b
Imiquimod - - + + + +
MCC950 - - - - + +
I
IL-1β release (pg ml-1)
0
500
1000
1500
2000
mCh Arl8b mCh Arl8b mCh Arl8b
Imiquimod - - + + + +
MCC950 - - - - + +
J
LDH release (%total)
0
20
40
60
80
100
mCh Arl8b mCh Arl8b mCh Arl8b
dA:dT - - + + + +
[8]Calix - - - - + +
K
Caspase-1 activity (vs vehicle)
0
2
4
6
mCh Arl8b mCh Arl8b mCh Arl8b
dA:dT - - + + + +
[8]Calix - - - - + +
L
IL-1β release (pg ml-1)
0
200
400
600
800
1000
1200
1400
mCh Arl8b mCh Arl8b mCh Arl8b
dA:dT - - + + + +
[8]Calix - - - - + +

**Supp. Figure 3. Effect of peripheral lysosome positioning on NLRP3 inflammasome activation in response to different NLRP3-activating stimuli, as well as AIM2 inflammasome activation, in iBMDMs. [A-I]** To assess NLRP3 inflammasome activation, iBMDM mCherry and iBMDM Arl8b-mCherry cells were primed with LPS (1 μg $mL^{-1}$) in complete media for 2 h. Following pre-treatment with PBS or MCC950 (10 μM) for 15 min, iBMDMs were stimulated with either silica (300 μg $mL^{-1}$) for 4 h **[A-C]**, ATP (5 mM) for 1 h **[D-F]** or imiquimod (Imiq; 150 μM) for 2 h **[G-I]**. Supernatants were then assessed for LDH release, caspase-1 activity and IL-1β release. **[J-L]** To assess AIM2 inflammasome activation, iBMDM mCherry and iBMDM Arl8b-mCherry cells were primed with LPS (1 μg $mL^{-1}$) in complete media for 2 h. Following pretreatment with DMSO or 4-Sulfonic calix[8]arene (10 μM) for 15 min, iBMDMs were activated by lipofectamine-mediated transfection of mock (lipofectamine alone) or poly(dA:dT) (1 μg $mL^{-1}$) for 4 h. Supernatants were then assessed for LDH release **[J]**, caspase-1 activity **[K]** and IL-1β release **[L]**. (n = 5 independent experiments). Data were analysed using two-way ANOVA with Holm-Sidak's multiple comparisons test for parametric data or unpaired two-tailed Mann-Whitney test for non-parametric data. ****$P \leq 0.0001$.
